# Mechanism of spike opening in a mammalian gammacoronavirus

**DOI:** 10.64898/2026.09.30.755498

**Authors:** John D. Clarke, Michalina Grabias, Antonio Di Maio, Harley Phlora, Snezana Vasiljevic, Sean A. Burnap, Rupa Tripathy, Wengang Chai, Weston B. Struwe, Donald J. Benton, Yan Liu, Steve J. Gamblin, Antoni G. Wrobel

## Abstract

Coronavirus spike proteins mediate receptor binding and fusion and are key determinants of viral host range. To bind protein receptors, some CoV spikes undergo conformational changes termed opening, which expose the receptor-binding surface. Yet, the prerequisites and mechanism of spike opening remain poorly understood, especially outside of beta-CoVs, and many CoV spikes have been observed only in the closed conformation. The receptor specificities of gamma-CoVs, for which birds are natural hosts, await in-depth characterisation and their spikes have not been observed to open. Here, we report structures of the spike from an unusual mammalian gamma-CoV isolated from a beluga whale (BwCoV), in closed and open conformations, the latter with two S1B domains erect. BwCoV spike opening is associated with changes throughout S1 that resemble those observed in beta-CoV spikes and similarly propagate into destabilisation of the fusogenic S2 core. Furthermore, the BwCoV-S structure contains an additional density likely corresponding to a fatty acid, consistent with lipid binding stabilising the closed conformation through a pocket distinct from that identified for beta-CoVs. These structural insights reveal conserved principles regulating spike dynamics across coronavirus genera and support lipid binding as a potential modulatory mechanism of spike opening.

## Introduction

The *Orthocoronavirinae*, a subfamily of true coronaviruses, comprises four genera: alpha- and beta-CoVs, which typically persist in mammalian reservoirs, and gamma- and delta-CoVs, for which the natural hosts are birds. Coronaviruses have repeatedly crossed host-species barriers, most recently exemplified by the zoonotic emergence of SARS-CoV-2 which led to the COVID-19 pandemic^1^.

The CoV spike protein is a virion-membrane-embedded type I fusion glycoprotein and a major determinant of viral host range. CoV spikes are homotrimers functionally divided into two subunits: the N-terminal, membrane-distal S1 subunit—typically comprising four domains S1A-D (Fig. 1A)—which engages host cell receptors, and a C-terminal, membrane-proximal S2 subunit, which mediates membrane fusion^2^.

**Figure 1.**
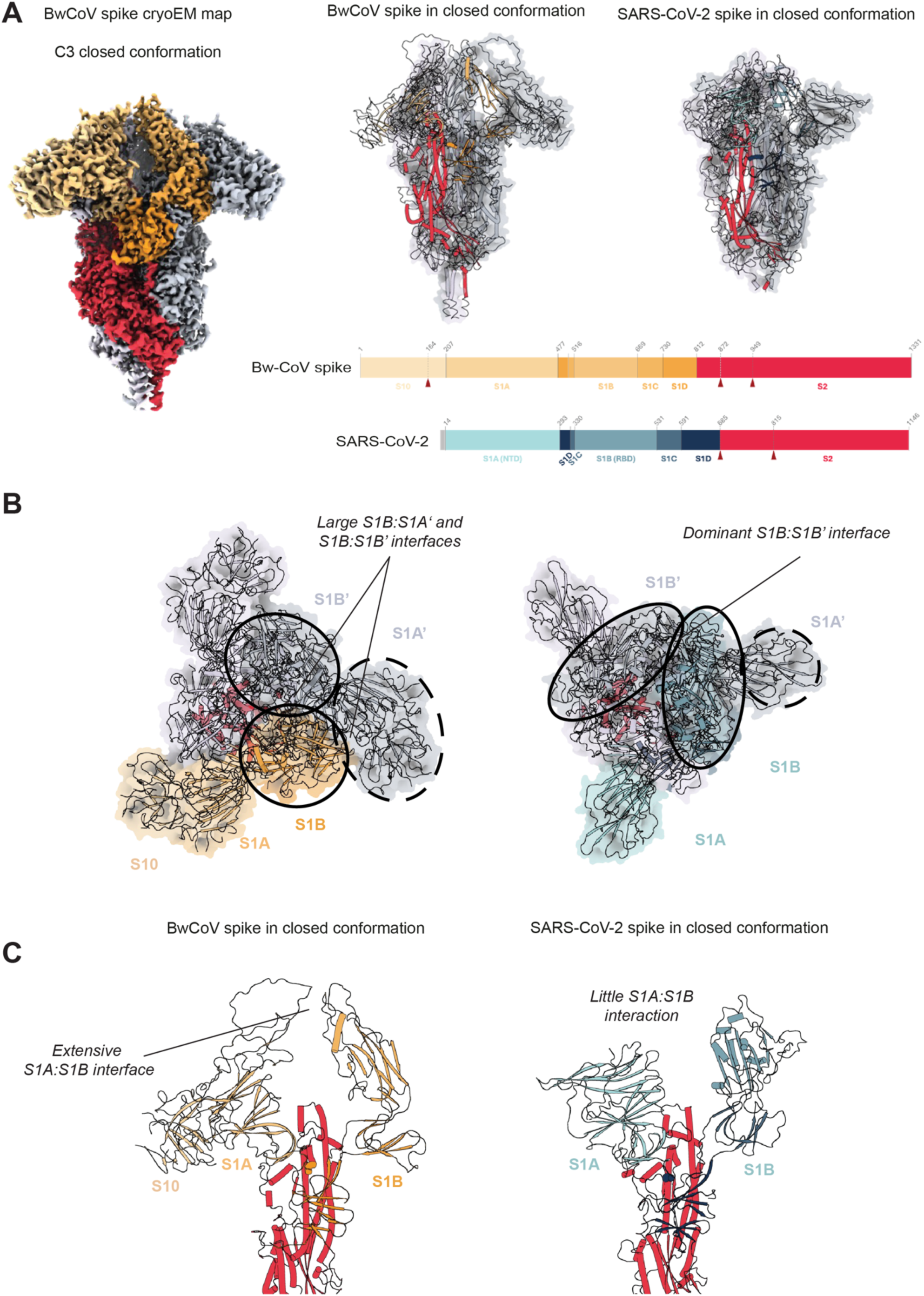
Closed conformation of BwCoV spike. (A) (left) C3 cryoEM map of BwCoV-S in the closed conformation and (top middle) the cartoon showing the corresponding structure compared to (top right) the closed conformation of a prototypical beta-CoV spike of SARS-CoV-2, PDB ID 6zge^5^. Domains are coloured as per the domain boundary schematic below, with progressively darker orange for the S1 subunit of Beluga-CoV-s (with an additional S10 domain compared to SARS-CoV-2) and cyan for the S1 of SARS-CoV-2 spike, and red for the S2 subunits of both. (B) Orthogonal views of the BwCoV-S (left) and SARS-CoV-2 spike (right) trimers, showing different spatial arrangement of the S1 domains stabilising the two closed conformations. (C) A single-protomer view of (left) the BwCoV-S and (right) SARS-CoV-2 in cartoon representation highlighting the intraprotomer interactions and additional features of the S1 of BwCoV-S. Colouring in all panels matches that in (A). All structures are aligned on the S2 connecting helix of chain A (residues 1125-1174).

CoV spikes recognise host glycans and proteins as receptors and undergo conformational rearrangements, which regulate access to their receptor-binding surfaces and drive membrane fusion. In some beta-CoVs that primarily use protein receptors—such as SARS-CoV, SARS-CoV-2, MERS-CoV, and PHEV—receptor-binding domains (RBDs, classified more generally as S1B domains) spontaneously sample exposed ‘open’ conformations, even in the absence of receptor^3–8^. Recently, spontaneous spike opening to present the putative receptor-binding motif has also been observed for the spike of PEDV, an alpha-CoV^9^. In contrast, spikes of several other coronaviruses have been observed to open only upon receptor engagement. This includes spikes of alpha-CoVs, 229E and FCoV, which open in the presence of their protein receptors^10,11^, and a beta-CoV, HKU1, in which binding a sialoglycan coreceptor to the S1A domain allosterically drives opening of the S1B to allow subsequent engagement of the protein receptor^12^. Other CoV spikes have been observed only in closed conformations despite using protein receptors, as exemplified by several bat CoVs such as RaTG13^5,13–15^. These observations suggest that spike opening can be either intrinsically sampled or induced by receptor presence, but the structural determinants that drive this process remain incompletely understood. In particular, no spike from the predominantly avian genera of delta- and gamma-CoVs has previously been observed in an open conformation.

The receptor engagement by gamma-CoVs has yet to be characterised at the molecular level. The most studied gamma-CoV, Infectious Bronchitis Virus (IBV), causes a severe respiratory disease affecting poultry industries globally^16^. Sialoglycans have been identified as potential IBV receptors^17^ but the mechanistic details of IBV entry remain poorly understood. Infections with gamma-CoVs have also been observed in mammals. BwCoV, found in a liver of a captive beluga whale (*Delphinapterus leucas)*, was the first gammacoronavirus discovered in a mammalian host^18^, and a related gammacoronavirus was subsequently identified in dolphins (*Tursiops truncatus*)^19^.

While this work was being finalised, structures of cetacean gamma-CoV spikes were published^20^ revealing an unusual S1 architecture, but little remains known about their receptors. Moreover, similarly to the IBV spike^21^, the Beluga spike has only been observed in one state, resembling the closed conformation of alpha- and beta-CoV spikes^21^ with all three S1B domains capping the fusogenic core, leaving unresolved whether gamma-CoV spikes can undergo opening and what structural features are associated with this transition. Here, we report structures of the BwCoV-S in both open and closed conformations, reveal potential determinants of the close-to-open transition, and take initial steps toward understanding BwCoV-S receptor recognition.

## Results

### Overall structure of the BwCoV spike

We determined the structure of the closed BwCoV-S by cryogenic electron microscopy at 2.8 Å resolution with single-particle analysis and C3 symmetry imposed to increase resolution (Fig 1A). The spike adopts a canonical homotrimeric quaternary topology for coronavirus spikes, with the addition of a 207-residue, N-terminal S10 domain upstream of the S1A domain within the same protomer, with which S10 makes primary interactions (Fig. 1B). The membrane-proximal part of the S1A resembles that of the beta-CoV spikes, with similar arrangements of S1A beta-sheets contacting the helix and beta strands, which cap the S1D and S1C domains (residues 294–330 in SARS-CoV-2 spike and 477-515 in BwCoV) and have been shown to be key contributors to the opening of beta-CoV spikes^22–25^. Furthermore, the ‘top’ (viral-membrane distal) part of the S1A extends further away from the membrane towards the vertex of the spike protein, forming a prominent long loop (residues 355-380). This loop contacts the 635-640 loop of the S1B domain of the same protomer, burying 170 Å^2^. This arrangement differs from beta-CoV spikes, such as SARS-CoV-2, in which there is little intra-protomer interaction at the viral-membrane-distal spike vertex (Fig. 1C). Inspecting the corresponding C1 reconstruction of the closed conformation reveals that this intra-protomer S1A:S1B interaction is lost for one chain: despite all three S1B domains remaining in the closed arrangement covering the S2 core, one of the S1A domains retracts away from the symmetry axis, breaking the interaction between its loop and the S1B of the same chain (Fig. S1A). This rearrangement reduces one of the three protomer:protomer interfaces by ∼400 Å², predominantly through a loss of an interprotomer S1A:S1B’ interface, while the S1B domains remain closed and the combined S2:S2 interface unchanged (Table S1).

In both C1 and C3 closed conformations, the S1A and S1B domains are arranged around the trifold symmetry axis, with both domains making extensive interactions with the adjacent protomers. In the C3 reconstruction, the S1A:S1B’ interface buries 550 Å^2^, while the S1B interacts with both adjacent protomers burying 130 Å^2^ with S1B’, 130 Å^2^ with S1B’’, and 550 Å^2^ with S1A’’ (Table S1). This makes the interactions between S1B and S1A domains (of both the same and adjacent protomer) key for maintaining the S1 trimer, unlike in beta-CoVs, in which the S1:S1 interactions between adjacent protomers are predominantly maintained by neighbouring S1B domains (Fig. 1B). The S2 domain of BwCoV resembles that of other CoVs and, as in them, makes extensive interactions with the neighbouring protomers and is capped by the S1 subunits. Unlike many beta-CoVs known to open, such as SARS-CoV-2 and MERS-CoV, BwCoV spike does not have an obvious furin site at the S1/S2 boundary. However, there are two potential polybasic cleavage sites within S2 (869RCRR872 and 947RDAR950), as well as a putative site within the S10 domain (162RHER165), all of which in our structure show good Coulomb-potential density and thus no indication of cleavage.

### BwCoV spike adopts closed and open conformations

The majority (74.5%) of BwCoV-S particles collected presented a ‘closed’ conformation, with the S1B domains oriented towards the trifold symmetry axis, and covering the S2 fusogenic core. However, unlike in the structures of other gamma- and delta-CoV spikes, the remainder of BwCoV-S particles were classified into an ‘open’ state (Fig. 2A), with two out of three protomers adopting a new conformation with the S1B domain uncovering the S2 core by rotating outward by approximately 39° compared to the closed state, and their centres of mass moving by 19 Å (Fig. 2B). The third S1B remains down but is rotated by ∼10° relative to S1B in the fully closed conformation, thus adopting an intermediate ‘ajar’ position between the closed and fully erect arrangement (Fig. S1B). As in other spikes which demonstrate an open conformation, the pivot point for the S1B rotation is located on the S1B:S1C linker, close to the S1B domain. As a result, the S1C remains largely in the same position upon the opening (Fig 2B). The local resolution of the map at the membrane-distal part of the structure is lower than in the closed conformation, consistent with increased flexibility of the S1B in both closed and open protomers (Fig. 2A). The outward (away from the symmetry axis) movement of the S1B is accompanied by outward and downward (towards the viral membrane) movement of the S10+S1A ensemble by ∼14° and a centre of mass translation of 6 Å (Fig. 2C), in line with similar movements observed for the S1A upon opening in beta-CoVs^5,24,26^. In addition, S10 pivots a further 3° towards the viral membrane (Fig. S1A). Notably, a similar repositioning of the S10+S1A ensemble is observed in all three protomers (14.4°, 14.1°, and 14.2°), including that with the down (‘ajar’) S1B, which furthermore also loses its interface with the prominent 355-380 loop of S1A. Together with the similar S10+S1A repositioning observed for one of three chains in the C1 closed reconstruction (Fig. S1A), this demonstrates that rearrangements within the S10+S1A ensemble and release of the apical S1A:S1B interaction can precede full opening of S1B.

**Figure 2:**
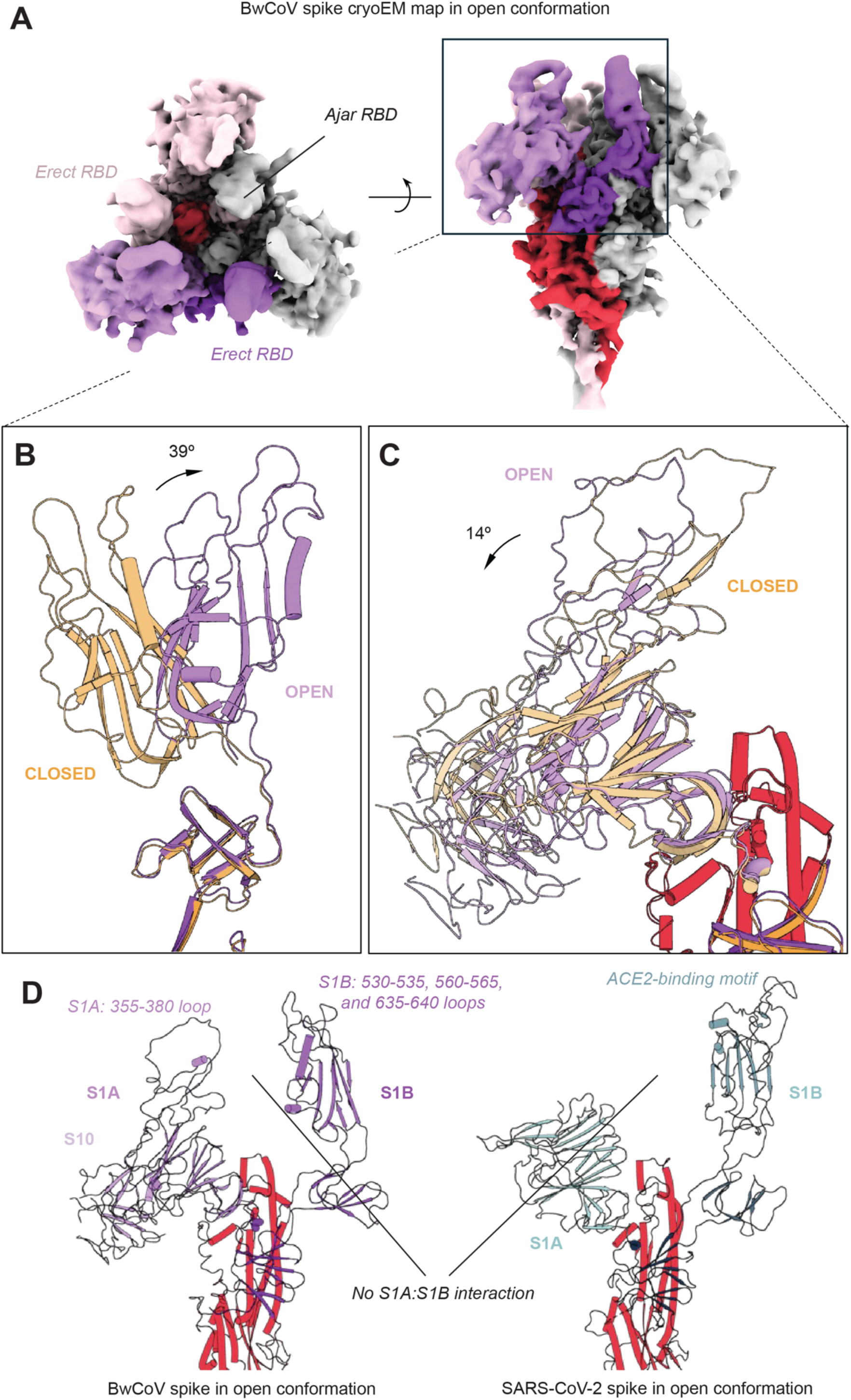
Inherent flexibility of the S1 subunit of BwCoV-S. (A) Orthogonal views of the cryo-EM map of the BwCoV spike ectodomain in the open conformation at a threshold of 0.350 showing two protomers in the open conformation and one in the partially-closed (‘ajar’) conformation. Colouring of S1 in the open structure matches that in Fig. 1A for the closed conformation but with purple instead of orange. (B, C) Motion of (B) the two fully erect S1B domains and (C) all three S1A domains upon opening. More minor repositioning is also observed for the third S1B and in the S10 domains (see Fig. S1). Colouring matches Fig. 1 for the closed conformation and panel 2A for the open. Structures have been aligned on the S2 connecting helix of chain A (residues 1125-1174). (D) Comparison between single opening protomer of (left) the BwCoV-S reported here and (right) SARS-CoV-2 in an open conformation (one-RBD-erect, PDB ID 6ZGG^5^) shown in cartoon representation.

### Characteristics of the open conformation of BwCoV spike

The closed-to-open transition limits the protomer:protomer interactions holding the trimer together, with the ∼6100 Å² buried interface between equivalent protomers in the closed C3 state reduced to between ∼4780 and 5130 Å² across the three non-equivalent protomer interfaces in the open state, and exposes new surfaces (Fig. 2D). Upon spike opening, both inter- and intra-protomer interfaces are affected. The interfaces between neighbouring S1B domains are completely lost in the open structure (∼130 Å² each), while the interactions between S1B and the S10+S1A domains of the adjacent protomer are substantially reduced or lost (∼460–550 Å²) (Fig. 3A). The interfaces between the two erect S1Bs and adjacent S2 domains are also lost (∼240 Å² each), but the corresponding interface involving the ajar S1B is largely preserved (215 Å² compared with 242 Å² in the closed structure) (Table S1). As seen for other type-I fusion proteins, these changes uncover the core central helices of the fusion apparatus, making space for them to extend towards the cell membrane during fusion. They are also associated with rearrangement within the S2 fusogenic core itself (Figure S1D): slight dilation, also observed in other type-I fusion proteins^22,27,28^, is accompanied by a combined loss of ∼520 Å² (∼7%) of buried surface area between the three S2 domains. The opening also changes the landscape of interactions within a single protomer: it exposes to solvent the prominent S1A loop (residues 355-380), which S1B contacts in the closed conformation, and the loops 530-535, 560-565, and 635-640 of S1B (Fig. 2D). In the two protomers with erect S1Bs, the S1B:S1C interface decreases by ∼170–180 Å² compared with the closed conformation, while it remains largely preserved in the protomer with ajar S1B (a decrease of only 11 Å²) (Table S1). In contrast, the ∼170 Å² intra-protomer interface between the S1A loop and S1B is lost in all three protomers. At the same time, rearrangements of S1A, S1B and the smaller S1C and S1D subdomains redistribute interactions throughout S1 (Table S1). This is reflected by changes in the interactions of S1C with adjacent S10+S1A and S2 domains, and increased interactions between S1A and S2 (+22–43 Å²) within the three protomers. S1D moves away from the S2 core, losing 48–94 Å² of interface within the same protomer, while its interaction with S1A remains largely preserved. The additional pivoting movement of S10 relative to S1A (Fig. S1C) is accompanied by substantial remodelling of their interface, which decreases by ∼110–310 Å² across the three protomers (Table S1). These changes identify the smaller subdomains S1C and S1D as key contributors to accommodating the open state, as extensively described for beta-CoVs, especially SARS-CoV-2^22–25,27^, and reveal how the S10 domain, absent in beta-CoV spikes, may further contribute to this process.

**Figure 3.**
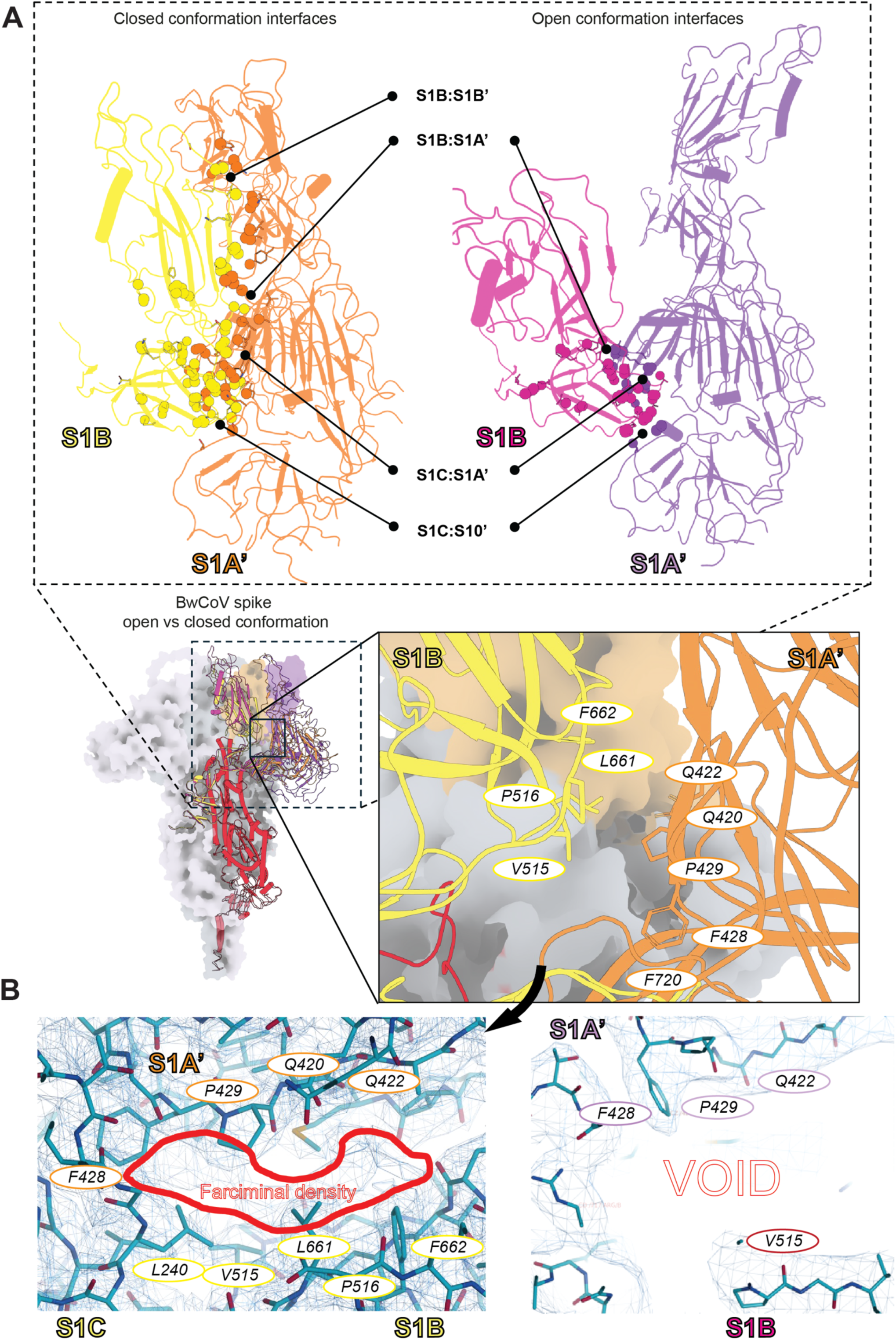
Characteristics of the open state of BwCoV-S. (A) Inter-chain interfaces of the Beluga Whale coronavirus spike S1 domain in (left) the C3 closed and (right) the open conformations. Each protomer is represented in block colours: in (left) the closed conformation in orange and yellow, and in (right) the open conformation in purple and pink. Residues that participate in the inter-domain interface are shown with stick notation, and interfacing atoms are shown as spheres. (Middle insets) (Left) Overall BwCoV-S open vs closed conformation coloured as in A, with boxes to show orientation for the remaining panels: panels in (A) share the same orientation, whilst those in (B) are rotated for clarity of showing the pocket between S1B and S1A. (B) (left) The pocket between S1B and S1A’ surrounds an additional farciminal density in the C3 closed conformation, (right) absent in the open conformation. Maps are shown at threshold 1.000 (see Figure S2 for the closed C1 map and remaining protomers).

The open conformation also lacks a large farciminal stretch of additional density between the S1A’ and the adjacent S1B domain, which we noticed while building the model of the C3 closed conformation but could not assign to any obvious protein or glycan component (Fig. 3B and S2A). This density is reminiscent of the free fatty acid binding pocket first observed in the Wuhan variant of the SARS-CoV-2 spike^5,29^, and later in spikes of SARS-CoV, MERS-CoV, and SARS-CoV-2 variants of concern^30^. The pocket where this density resides is predominantly hydrophobic—lined by residues such as F428 of S1A and V515, F662, F720 of the adjacent S1B’—but capped by hydrophilic Q420 and Q422 (S1A), which could engage the carboxyl group of a fatty acid (Fig. 3B). Indeed, modelling of stearic acid into the density reveals that it could accommodate 18-carbon-chain fatty acids such as stearic, oleic, or linoleic acid (Fig. S2B). In the published structure of BwCoV-S^20^, this density is present but rather diffuse (Figure S2A), and in our closed C1 map it is prominent in two protomers but substantially weaker in the pocket lined by the S1A that resides slightly away from the symmetry axis, consistent with ligand occupancy favouring the closed conformation (Figure S2A).

### Towards understanding the receptor binding by BwCoV spike

In the CoV genera for which the spike opening has been observed, the transition to the open conformation is normally a prerequisite for binding to a protein receptor. The most common protein receptors of CoVs include angiotensin converting enzyme-2 (ACE2) aminopeptidase N (APN), and dipeptidyl peptidase-4 (DPP4), although other receptors such as dipeptidase1 or CEACAM1a have also been described^7,31^. We produced the most common putative receptors as recombinant ectodomains of ACE2, APN, and DPP4 from beluga whale and tested their binding to BwCoV-S using biolayer interferometry and mass photometry. However, none of these proteins showed convincing evidence of spike binding at the < 1μM concentrations tested (Fig. S3).

Glycans are another common class of CoV receptors, shown not only to directly bind to CoV spikes^32–34^ but also to coordinate engagement of the protein receptors^12^. Sialosides have been also the only receptors of the gamma-CoVs identified to date with especially α2,3-sialylated glycans showing prominent binding to IBV in glycan arrays^17^. We thus screened BwCoV-S against a focused glycan microarray comprising 40 glycans selected from our larger, recently published glycan library^35^, spanning major structural classes of the mammalian glycome (Table S2). BwCoV-S showed response to sulphated glycosaminoglycan (GAG) probes, particularly heparin hexa- and deca-saccharides, but also chondroitin sulphate probes. Weak but detectable binding was observed to long chain sialyl glycan probes 6’Sia-Octa-T2 and Fetuin N-Glycan A2 (Fig. 4). This specificity is partially reminiscent of SARS-CoV-2 spike, which was shown to bind unmodified N-acetylneuraminic acid and α2,3- and α 2-6-linked sialosides^33^. This reveals glycans as spike interactors and thus potential receptors or attachment factors of BwCoV.

**Figure 4.**
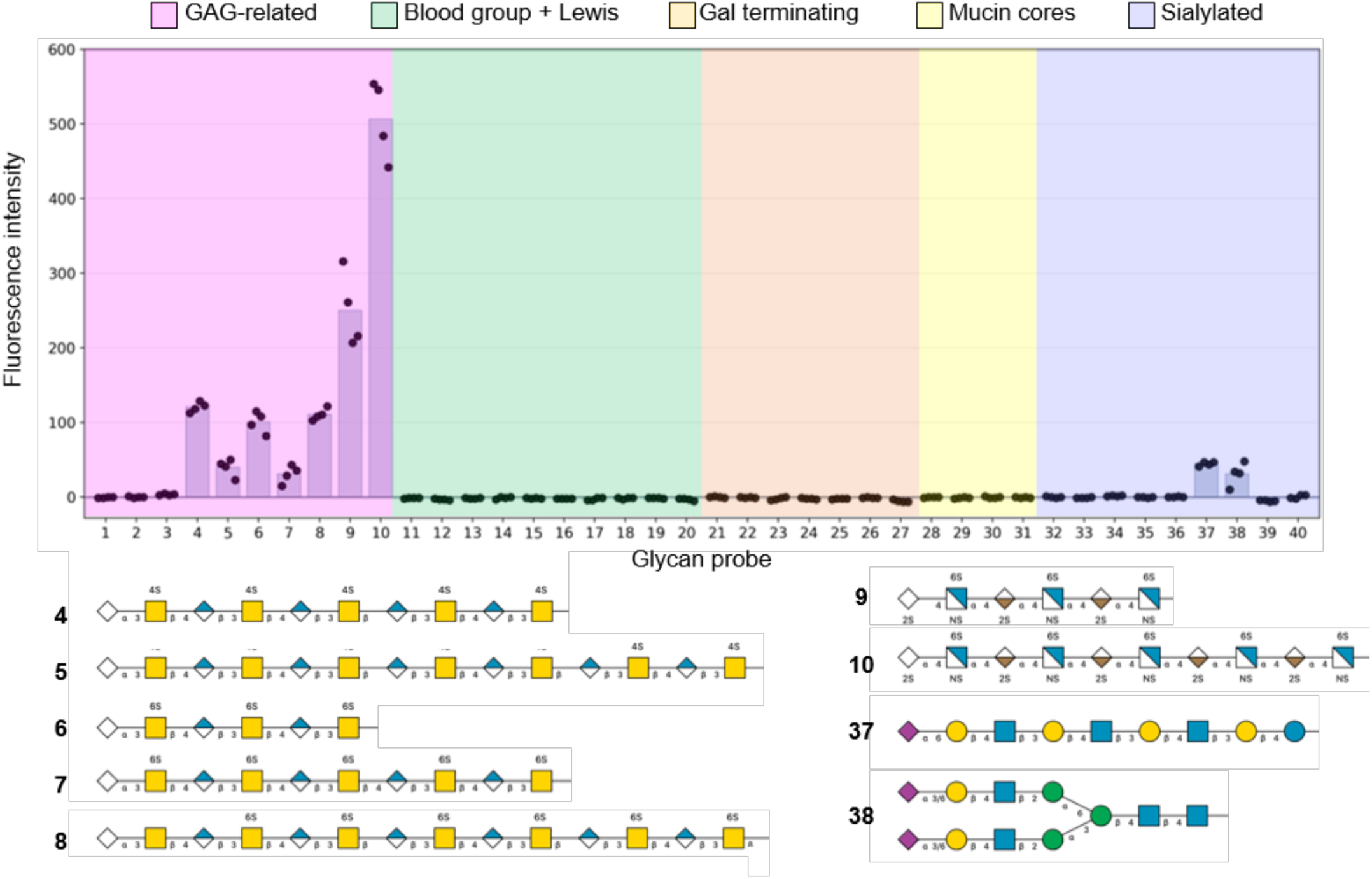
BwCoV-S binding to glycans on a focused microarray. Bar chart showing the mean fluorescence intensities for BwCoV-S binding to individual glycan probes. Results are shown as the mean of quadruplicate spots. Structures of the glycan probes showing a positive response with BwCoV-S, depicted using Symbol Nomenclature for Glycans (SNFG). The sequences of all glycan probes on the microarray are provided in Table S2.

## Discussion

Here, we report structures of a gamma-CoV spike in the closed and open conformations—the latter with two S1B domains erect. This open structure of BwCoV is unusual, as all previously reported structures of related gamma- and delta-CoVs have only been observed in the closed conformation. Compared to beta-CoVs, for which the opening is best understood, the BwCoV-S opening shows differences, while also sharing key similarities. Contrary to beta-CoVs, BwCoV spike shows considerable interactions between the S1A and S1B of the same protomer at the spike’s vertex, which are lost upon opening. To accommodate this, the S1A moves outward and downward as S1B rotates out while the smaller S1C and S1D subdomains rearrange their interactions during the closed-to-open transition. Notably, in the closed C1 structure, the asymmetric S10+S1A rearrangement is accompanied by loss of both intra- and inter-protomer S1A:S1B contacts while all S1Bs and the S2 core remain in their closed arrangement. A similar rearrangement is retained in the S1B-down protomer of the open structure, in which S10+S1A undergoes a similar ∼14° rotation to that in the two protomers with erect S1Bs, while S1B itself adopts an ‘ajar’ orientation rotated by only ∼10°. These observations suggest that remodelling of the S1A:S1B interface and movement of S10+S1A may precede bringing up the S1B domain, consistent with a sequential rather than concerted opening mechanism. A remodelling of another S1A:S1B interface has been reported before to affect the opening of HKU1 spike upon glycan binding^12^. This may suggest that the S1A-to-S1B coupling is a conserved mechanistic principle controlling exposure of S1B across very divergent coronavirus spikes.

As with other spikes, the BwCoV-S trimer opening leads to reduced inter-protomer interactions, a loss of ∼1000–1300 Å² of buried surface area between protomers, thereby destabilising the trimer. This is, however, still much less than when a prototypical protein-binding CoV spike of SARS-CoV-2 opens, which loses >2000 A^2^ to bring up only one S1B^27^, despite a comparable protomer interface of ∼ 6000 A^2^ in both closed trimers. However, similar to SARS-CoV-2 spike^27,28^, the rearrangements in S1 upon BwCoV-S opening destabilise the S2 core through a combined loss of ∼520 Å² of its trimerization interface, consistent with priming the spike for fusion.

Another structure of BwCoV-S (PDB ID: 9R1Q) has been reported recently by Hulswit et al., while this work was in its final phase^20^. This study provides remarkable insights into BwCoV biology identifying the S10 domain as a member of the cupin protein superfamily and providing detailed insights into BwCoV-S glycosylation. Notably though, the authors have not observed the BwCoV-S in an open conformation: they define the S1B domains as upright, however these remain capping the S2 core as in our closed conformation. This could be due to minor differences in the protocols used throughout the study: during protein production, cryoEM sample preparation, or data processing. Such differences are, however, not unprecedented for CoV spikes as especially structures of SARS-CoV-2 have been observed in multiple very similar conformations, despite almost identical construct design and production conditions^4,5,24,29,36^. A detailed comparison of 9R1Q with our closed conformation (Figure S4A) reveals that this might be the case here. For instance, a small repositioning (a 3° rotation outward, with respect to the threefold symmetry axis) of the S10 is observed in the way the S10 abuts S1A, providing a clue to the flexibility of the BwCoV spike (Figure S4B). This movement appears to be a rigid-body motion, as the S10 domains can be superimposed with those of Hulswit *et al.* with an RMSD of 0.9 Å (Figure S4C). Indeed, each successive domain of BwCoV-S is very similar to the earlier structure, save the C terminus of the S2 domain, which demonstrates an average elongation of ∼1 Å (Figure S4A), most pronounced at the stem helix (∼1.5 Å). Notably, these differences between the previously reported closed conformation and the one we report here are along the same axes as those we observe comparing our closed and open conformation. Moreover, our closed C1 reconstruction already shows loss of the apical S1A:S1B interaction in one protomer while all three S1B domains remain in the closed orientation, providing further evidence of conformational variability within closed populations. This suggests that 9R1Q may thus represent a slightly more closed or compact conformation of the BwCoV-S than the one we observed, which could contribute to why an open class was not observed, and be a similar, albeit less extreme, case of the closed vs locked conformations observed for SARS-CoV-2 spike^36^. This could also be due to the differences in the unassigned farciminal density observed between the neighbouring S1A and S1B’ domain in the map reported by Hulswit et al. and in our closed structure that likely corresponds to a lipid. Together, these observations suggest that 9R1Q, our C1 closed structure, and our open structure may sample a conformational continuum.

Spikes of beta-CoVs have been shown before to bind free fatty acids^30^ and the additional density that we noticed in our closed BwCoV-S may suggest that this ability may extend to other CoV genera. For beta-CoVs a fatty acid, linoleic acid, has been shown to stabilise the spike trimer in a ‘locked’ prefusion conformation with S1B domains restricted in movement,^37^. Furthermore, linoleic acid reduced virus infectivity possibly by restricting spikes’ ability to open^29,37,38^. A homologous pocket is conserved in various beta-CoV spikes, including those of SARS-CoV, MERS-CoV, SARS-CoV-2, HCoV-OC43, and HCoV-HKU1 (though in some of them the entrance to the pocket is occluded, as, e.g., HCoV-HKU1 cannot bind linoleic acid)^30^ suggesting its functional significance in beta-coronaviruses. The unexplained density consistent with a free fatty acid that we observed in the closed conformation of BwCoV-S but not in the open conformation suggests that lipid binding may represent a more general mechanism regulating spike opening across CoV genera. Consistent with this, the density is prominent in two protomers of our C1 closed reconstruction but substantially weaker in the pocket associated with the S1A shifted away from the symmetry axis and of reduced interface with the neighbouring S1B’. Remarkably, the putative pocket in our structure is located on a different interface from that observed before: in beta-CoV spikes, it resides between adjacent S1B domains, while we observe it between S1A and the adjacent S1B’. While the pocket in BwCoV-S does not show very strong conservation when compared with other gamma-CoVs (Fig. S5), its observation suggests convergent use of lipid binding to tune the closed/open equilibrium in CoV spikes through different structural solutions. It is conceivable that BwCoV-S acquired ability to open—along with that of binding lipids which may regulate it—during its adaptation to a mammalian host, especially as prototypical gamma-CoV spike of IBV has not been observed to open^21^. This could coincide with a receptor change required to establish in a new host.

The ability of BwCoV spike to open may provide clues about its receptor recognition. For many CoVs, S1B opening exposes a protein-receptor-binding surface. The two regions that become liberated upon opening include the large loop of S1A and several smaller surface loops of S1B. Analysis of the exposed surface electrostatics suggests that opening reveals a small negatively charged cavity on the S1B that could potentially engage a binding partner (Fig. S6). However, while protein receptors are most often bound by the S1B domain in other spikes (except MHV^39^), the unique interaction between S1A and S1B of the same protomer at the membrane-distal part of the spike, as well as the prominence of the S1A loop, may suggest that receptor binding in Beluga-CoV evolved differently, with S1A also playing a direct role in the receptor-engagement. The identity of the potential protein receptor remains to be elucidated though: while we did not observe binding of typical CoV receptors (APN, ACE2, and DPP4) to Beluga-CoV, they cannot be fully excluded, as various factors such as membrane context or glycosylation may regulate their binding, nor can any other protein receptor be ruled out.

Glycans are established CoV receptors and attachment molecules, engaged most often via binding pockets on the S1A domain^31–33^. Investigating the surface of BwCoV-S reveals exposed, positively charged clefts on the S1A and S10, which could accommodate negatively charged glycans, including sialosides and GAGs (Fig. SF6). Together with the glycan microarray results, this suggests that these glycans may serve as attachment factors for Beluga-CoV. While little is known about cetacean glycosylation, using sialosides and GAGs as receptors and attachment molecules is well described for other CoVs, including IBV^17^ and SARS-CoV-2^33,40^.

The unusual two-S1B-erect opening of BwCoV-S may relate structurally to the receptor used perhaps suggesting its oligomeric nature. It is also possible that, despite opening, the BwCoV spike doesn’t use a protein receptor and instead utilises this process to engage other receptors such as, e.g., two different types of glycans. In general, the ability of spike to bind glycans and the shared S1A-S1B remodelling as observed for HKU1, suggest shared general principles of receptor engagement among different spike genera. Finally, the remodelling and loss of interfaces within the S2 core suggest that the open conformation we observe is structurally primed for fusion. This might mean that the observed open structure has significance beyond receptor binding, perhaps representing a state on the way to fusion but without the ability to progress further given the 2P mutations used to stabilise the pre-fusion conformation here.

While further studies are required to reveal the details of receptor recognition by BwCoV, here we show that the S1A–S1B coupling with concomitant rearrangements in S1C and S1D may represent a mechanism controlling S1B exposure across divergent coronavirus spikes, that S1 opening propagates to the S2 fusiogenic core potentially priming it for fusion, and that beta-CoVs and BwCoV might have independently evolved lipid-sensitive structural features that favour closed spike conformations. Thus, divergent CoV spikes may share mechanistic principles of opening and fusion priming while achieving them through distinct molecular solutions.

## Acknowledgments

The authors thank Andrea Nans from the Crick Structural Biology Science Technology Platform for assistance with data collection; Andrew Purkiss, Andrea Nans (Crick Structural Biology Science Technology Platform), the Scientific Computing Science Technology Platform (Crick), and Ed Lowe and the COSMIC and computing teams (Oxford) for computational support; Valeria Calvaresi, Sophie Lawrence and members of Carol Robinson’s group, and the Wrobel lab for discussions. This work was supported by the Wellcome Career Development Award (303026/Z/23/Z) to A.G.W. and Early-Career Award (312750/Z/24/Z) to S.A.B., and by the European Union ERC grant GLYCvsPROT (101165048) to A.G.W. Views and opinions expressed are however those of the author(s) only and do not necessarily reflect those of the European Union or the European Research Council. Neither the European Union nor the granting authority can be held responsible for them. This work was also supported by the Francis Crick Institute which receives its core funding from Cancer Research UK (CC2060), the UK Medical Research Council (CC2060), and the Wellcome Trust (CC2060). The glycan microarray work was supported by the Wellcome Trust Biomedical Resource Grant (218304/Z/19/Z) and the March of Dimes Prematurity Research Centre (22-FY18-82) to Y.L.. Y.L. also thanks the members of the Glycosciences Laboratory, particularly Drs Yibing Zhang and Virginia Tajadura-Ortega, for their contributions to glycan probe and array preparation, and Professor Hongzhi Cao for providing glycan materials used in array construction. We also acknowledge funding from the UK Research and Innovation Future Leaders Fellowship to W.B.S. (MR/V02213X/1).

## Data availability

The EM models and maps have been deposited in the PDB^45–47^ and the EMDB^55^ as follows: BwCoV-S closed conformation with C3 symmetry imposed (PDB ID 34OG, EMD-100055), BwCoV-S C1 closed conformation (PDB ID 34SV, EMD-100140), and BwCoV-S C1 open conformation (PDB ID 34SW, EMD-100141).

## Authors’ contributions

D.B. and A.G.W. conceived the study. J.D.C., M.G., A.D.M., H.P., S.V., S.A.B., R.P., W.C., D.B., and A.G.W. performed the research. J.D.C., M.G., A.D.M., H.P., S.V., S.A.B., D.B., Y.L., and A.G.W. analysed the data. D.B., S.J.G. and A.G.W. supervised the structural biology work. Y.L. supervised and performed the glycan-array work. S.A.B., W.B.S., Y.L., S.J.G., and A.G.W. acquired funding. A.G.W. wrote the manuscript with contributions from all authors.

## Competing interests

The authors declare no competing interests.

## Materials and Methods

### Construct design

A construct containing the beluga whale coronavirus spike protein residues 1-1400 (GenBank ID YP_001876437) codon-optimised for human expression was synthesised and cloned into pcDNA3.1(+) by GenScript (UK), with an N-terminal μ-Phosphatase secretion leader sequence, and a C-terminal decahistidine tag for affinity chromatography. A dual-Proline substitution was made at positions 1148-1149 to stabilise the pre-fusion conformation^41^ and a TEV-cleavable FoldOn trimerisation domain was added between the C-terminus and the decahistidine tag. The putative beluga receptor constructs (ACE2 residues 19-614 (Uniprot ID: A0A2Y9M9H3), APN residues 67 – 969 (Uniprot ID: A0A2Y9MRE5), DPP4 residues 39 – 766 (Uniprot ID: A0A2Y9N2E9)) were codon-optimised for human expression, synthesised, and cloned individually into pcDNA3.1(+) by Genscript with an N-terminal Igκ light chain signal peptide and a C-terminal DYK-tag followed by a twin-strep tag (DYKDDDDKWSHPQFEKGGGSGGGSGGSSAWSHPQFEK).

### Protein expression

BwCoV-S was expressed in Expi293F cells (Gibco) cultured in suspension at 37 °C, shaking in 8% CO_2_ at 125 rpm. The cells were grown in the FreeStyle 293 Expression Medium and, at cell density of 3 × 10^6^/mL, 1 mg of the spike DNA was transfected per litre of the cell culture. The following day, the culture was enhanced following the manufacturer’s instructions and moved to 32°C, where it stayed for another three days before being harvested. The putative receptors were expressed in 293F cells without enhancement and remained for 6 days at 37 °C after transfection.

### Protein purification

To purify the protein of interest, the supernatant containing it was separated from the cells by centrifugation at 6000 RCF for 20 minutes. The supernatant was then filtered and 2x cOmplete™ protease inhibitor tablets (EDTA-free) (Roche) added together with either (for the spike) pre-washed (using Sample Buffer: 20 mM TRIS, 150 mM NaCl, pH 8.0) TALON beads (TAKARA; 10 mL bead slurry per 1 L culture supernatant) or (for the putative receptors) Strep-Tactin 4Flow resin (IBA Life Sciences). The mixtures were then incubated rolling for approximately two hours at room temperature before being applied to a gravity flow column. The resulting column was then either (for the spike) washed with four volumes of Sample Buffer, followed by four bed volumes of Wash Buffer (Sample Buffer + 20 mM Imidazole), and then eluted using eight bed volumes of Elution Buffer (Sample Buffer + 200 mM Imidazole) or (for the putative receptors) washed with eight bed volumes of buffer W (IBA Life Sciences) and eluted with three volumes of buffer BXT (IBA Life Sciences). The obtained flowthrough, wash, and eluted fractions were characterised by SDS-PAGE. Fractions containing proteins of expected size were pooled, spin concentrated, and injected to a Superdex™ 200 Increase 10/300 GL (Cytiva) column for Size Exclusion Chromatography in 20 mM TRIS, 150 mM NaCl, pH 8.0. Fractions of the eluate were characterised by SDS-PAGE, and those containing the protein of interest were pooled and spin-concentrated using appropriate Amicon® Ultra centrifugal filtration units (100 kDa MWCO for the spike and 30 kDa MWCO for the putative receptors).

### cryo-EM sample preparation and data acquisition for BwCoV-S

BwCoV-S supplemented with Octyl glucoside to a final concentration of 0.1 % (v/v) was applied to glow-discharged Quantifoil R 2/2 on 400 Cu mesh at 0.6 mg/mL using a Vitrobot mk3. Grids were glow discharged at 25 mA for 30 s and blotted for 4.5-5 s at 100% humidity and 4°C with pre-equilibrated filter papers, before being plunged into liquid ethane.

Two data sets were obtained from the same set of frozen grids: 1,810 movies were acquired on Talos Arctica operating at 200 kV with a Falcon 3 detector as a screening dataset, and then 10,224 movies were acquired using EPU 2.9 on a Titan Krios, operating at 300 kV, equipped with a Falcon 3 detector (Thermo Scientific) operating in the electron-counting mode. A total dose of 35 e−/Å^2^ was applied over 30 frames at a pixel size of 1.087 Å/px (on Titan Krios) and 43.7 e−/Å^2^ over 10 frames at a pixel size of 1.61 Å (on Talos Arctica) with a nominal defocorum range 0.5-2.5 μm.

### Single particle cryo-EM data processing of BwCoV-S

Movies were imported and motion- and CTF-corrected using RELION 3.1^42^. For both datasets, a subset of ∼1000 particles was manually picked with crYOLO^43^ to create a picking model (a new one for each dataset), and then all micrographs were picked using this model. Extracted particle stacks were 2D classified in RELION, and–for the screening dataset–they were exported to cryoSPARC^44^ to obtain an *ab-initio* 3D reconstruction. In the screening dataset the obtained *ab initio* model was then used for 3D classification with 15 classes in RELION^42^, and distinct 3D classes were identified corresponding to BwCoV-S in closed and open conformations, which were then refined in RELION. The obtained open conformation from the screening dataset was then used as a model for 3D classification in RELION for the main dataset resulting, again, in clear classes corresponding to open and closed conformations. These classes were then refined in RELION and particles corresponding to them Bayesian polished in RELION also. The resulting polished particle stacks were imported to cryoSPARC together with RELION models and homogeneously refined therein first in C1 and then, for the closed conformation, in C3 to increase resolution. The final structures were reconstructed from 172,370 for the closed and 59,112 for the open conformation. An overview of the processing workflow including final refinement FSC curves and representative densities is provided in Fig. S7 and Fig. S8.

To generate the closed structure, a published model BwCoV-S (PDB^45–47^ ID: 9R1Q^20^) was rough-body fitted to the resulting closed-conformation C3 map Gaussian filtered by 5 standard deviations using UCSF ChimeraX v1.11.1^48–50^ and iteratively fitted using Namdinator^51^ with decreasing stringency map filtering. Once satisfactory map fitting was achieved, the model was iteratively real space refined using Phenix v2.1-6048^52^ and manually rebuilt using Coot v1.1.20^53^. The obtained C3 closed model was then rigid-body fitted to the C1 closed and open maps, and iterative fitting by Namdinator^51^ and subsequent Phenix^52^ real space refinement and Coot^53^ manual rebuilding performed. For both models, once model building was complete, final statistics were determined using Phenix cryo-EM model validation (Table S3). Structural figures were generated using The PyMOL Molecular Graphics System v3.1.0 (Schrödinger LLC), and Coot^53^.

### Mass photometry

Mass photometry (MP) measurements were conducted using a Refeyn TwoMP system as previously described.^54^ High Precision No. 1.5H glass coverslips were cleaned via sonication in Milli-Q H_2_O, followed by isopropanol and Milli-Q H_2_0 then dried under nitrogen flow. Sample chambers were assembled using silicone gaskets (CultureWell™ reusable gasket, 3mm diameter x 1 mm depth, Grace Bio-Labs). Coverslips were placed on the sample stage and a single gasket was filled with 5-20 µL DPBS (without calcium, without magnesium, pH 7.4 ThermoFisher Scientific) to find focus and ensure low background signal-to-noise. Samples were measured at a final concentration between 10 and 50 nM. Acquisition settings within AcquireMP (v2023 R1.1, Refeyn Ltd) were as follows: regular field of view, frame binning = 2, frame rate = 498.3 Hz, pixel binning = 6, exposure time 1.95 ms and movies were taken over 60 seconds. Mass calibration was conducted using an in-house protein standard. Data were analysed using DiscoverMP (v2024 R2.1, Refeyn Ltd). Molecule counts were used to determine levels of protein complex formation. Representative histograms were generated using DiscoverMP.

### Biolayer interferometry

Biolayer interferometry was performed as established before for SARS-CoV-2 spike and ACE2^5,14^. The OctetRED384 system (Forte Bio) was used, and BwCoV-S immobilised on NTA Biosensors (Sartorius) at 100 nM, for 5 minutes, in 20 mM TRIS, 150 mM NaCl, pH 8.0 buffer. The immobilised sensors were then dipped in serial dilutions of the receptor solutions at 125 to 500 nM in the same buffer for 45 seconds to measure the association followed by a 2-minute dissociation step. All measurements were taken at 25°C.

### Glycan Microarray

The focused glycan array was prepared by printing 40 amino-linked onto high-density CodeLink slides (Surmodics) under conditions recently described^35,55^. The full list of glycan probes included in the array is provided in Table S2. To identify potential glycan ligands for BwCoV-S, purified His-tagged trimeric BwCoV-S was applied at 400 μg/mL to the array pre-blocked with 10 mM HEPES (pH 7.4), 150 mM NaCl, 1% (w/v) bovine serum albumin (Sigma 7030) and 0.01% Tween 20. Bound BwCoV-S was detected using a mouse monoclonal anti-polyhistidine primary antibody (Sigma-Aldrich SAB4200620), followed by a biotin-conjugated goat polyclonal anti-mouse IgG secondary antibody (Sigma B7264) and a 30-minute incubation with streptavidin–Alexa Fluor 647 (Molecular Probes) at 1 µg/mL. Microarray was scanned with a GenePix 4300A scanner (Molecular Devices) at 5 µm per pixel, using the 635 nm channel with a photomultiplier tube (PMT) setting of 400 and laser power of 100%. Images were analysed in GenePix Pro 7 with automatic spot alignment followed by manual adjustment. Mean fluorescence intensity after local background subtraction (Mean − B) was calculated from quadruplicate spots.

## Notes

### Competing Interest Statement

The authors have declared no competing interest.

