## Supplemental Information for "Mechanism of spike opening in a mammalian gammacoronavirus"

*Figures and Tables*

**Clarke et al.**

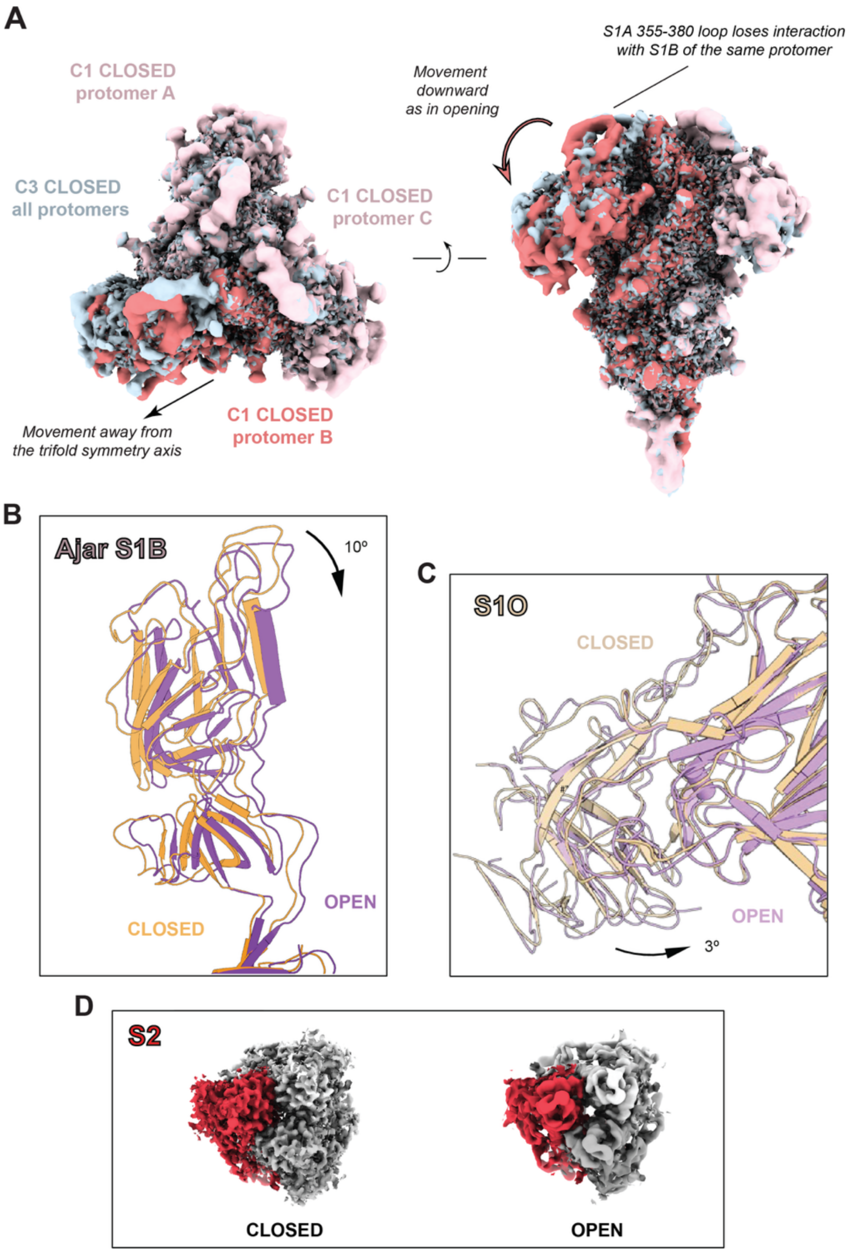


**Figure S1. Domain movements associated with Bw-CoV spike opening.** (A) Superposition of the BwCoV-S C3 closed reconstruction (blue) and the C1 closed reconstruction, with two protomers coloured pale pink and the asymmetric protomer coloured red (maps at contour 0.4). While all three S1B domains remain in the closed orientation, the S10+S1A ensemble of the red protomer in the C1 reconstruction moves away from the symmetry axis, disrupting both intra- and inter-protomer S1A:S1B interactions (Table S1). (B) Superposition of the S1B domains of the closed (yellow) and the chain assuming 'ajar' arrangement in the open conformation (purple): the ajar S1B is rotated partially away from S2. (C) Superposition of the closed (pale yellow) and open (pale violet) S10 domains: additional to the S1A movement, the S10 is further rotated towards the BwCoV-S S2 domain by 3°, when aligned on the S1A (residues 208:476). (D) CryoEM maps (coloured as in Fig. 1) in the top-down (towards the virion) view show S2 subunit in C3 closed (left) and open (right) conformations revealing dilation upon opening; the maps are at contour levels enclosing an equivalent volume (100,000 Å$^3$) to allow direct comparison between independently reconstructed maps with different density scaling.

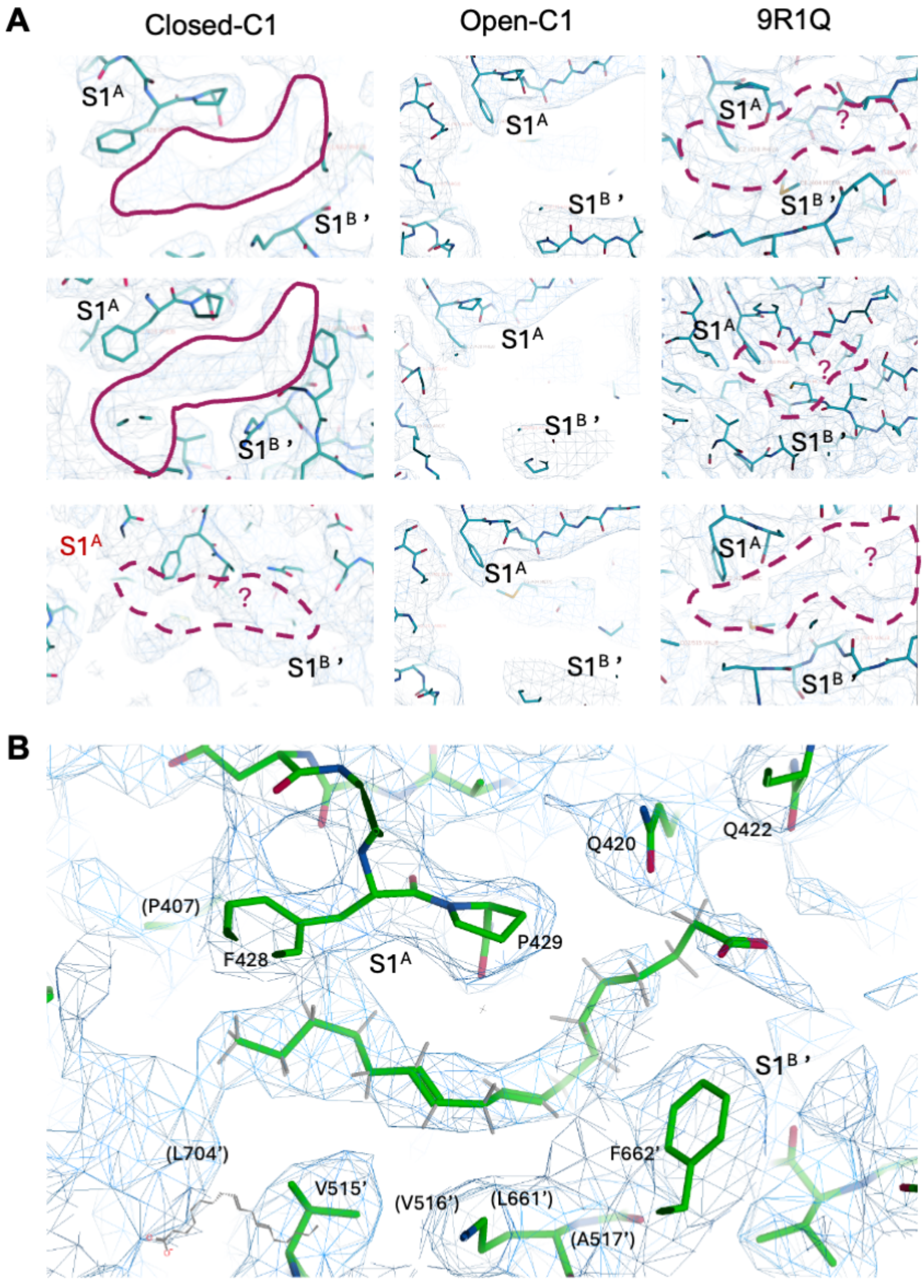


**Figure S2. Putative fatty-acid-binding site in BwCoV-S.** (A) Views on each trimer protomer (rows) showing unsatisfied farciminal density between the S1A' and the S1B of an adjacent protomer in the closed (left column) but not in the open (middle column) C1 non-symmetry imposed cryo-EM maps described in this study, compared to (right column) the published structure PDB ID: 9R1Q, EMDB-53512. Bottom left panel shows the pocket associated with the S1A (red label) which moves away from the symmetry axis, in which the corresponding density is substantially weaker than in the other two protomers. All maps visualised as blue mesh at a threshold of 1.000 in UCSF ChimeraX, with atomic models overlaid. Top middle same as in Fig. 3B. (B) Modelling of a C18 free fatty acid in the putative BwCoV-S closed conformation pocket revealing good fit to the density. Hydrophobic residues lining the pocket are labelled (with those in parentheses proximal to the pocket, but out of the imaging plane in the current view), as are Q420 and Q422 of the S1A which could cap the carboxylic acid group of a putative free fatty acid.

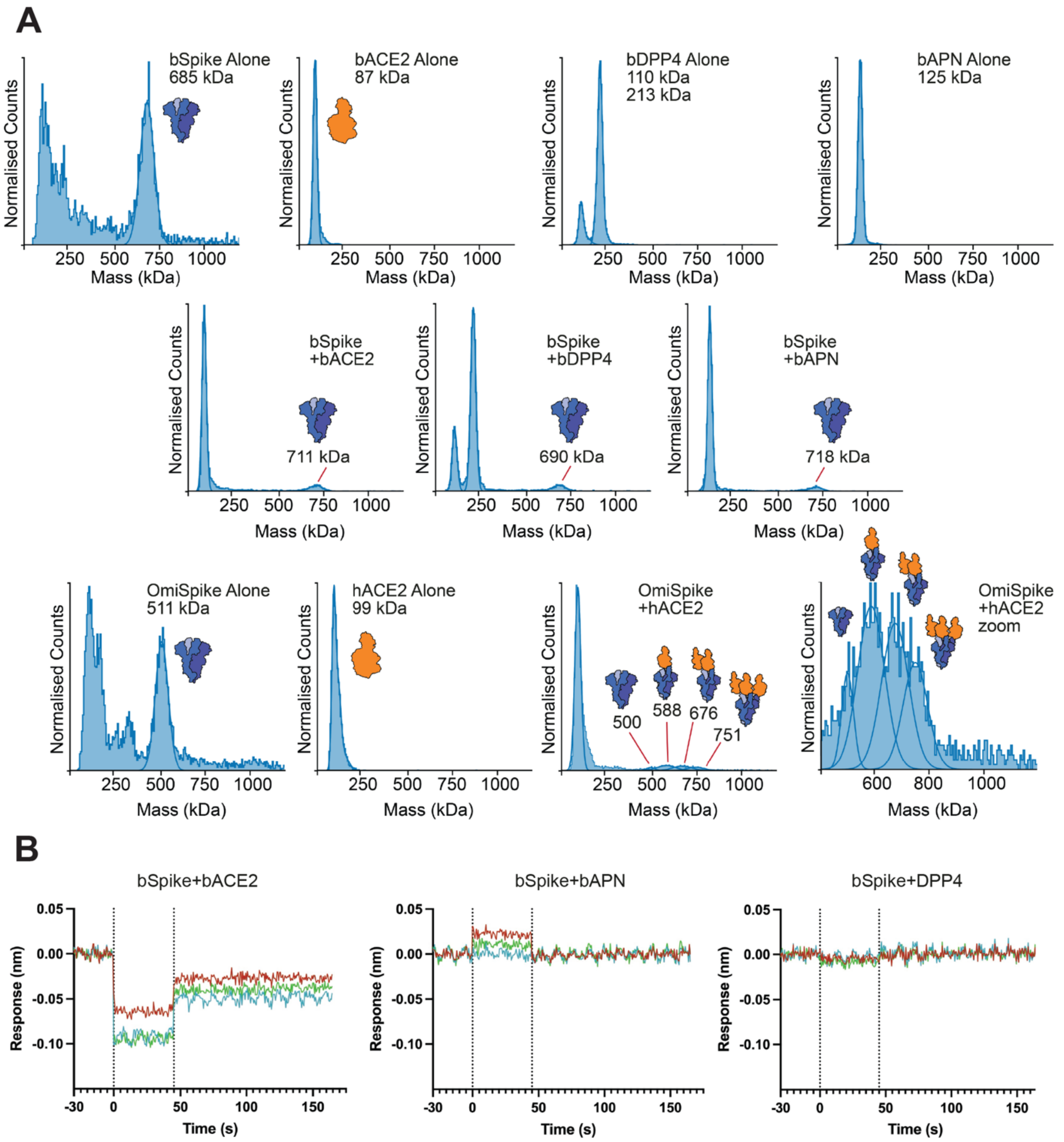


**Figure S3. Common protein receptors of CoVs do not bind to BwCoV-S.** (A) Mass photometry analysis of BwCoV-S binding to common CoV receptors. (top row) Representative MP mass histograms of BwCoV-S (bSpike) and candidate receptors from beluga: bACE2, bDPP4, and bAPN, measured alone at 20nM. (middle row) Representative MP mass histograms of the interaction between bSpike and candidate receptors bACE2, bDPP4, and bAPN at a 1:3 equimolar ratio (spike:receptor). (bottom row) A positive control: representative MP mass histograms of Omicron Spike (OmiSpike) and receptor human ACE2 (hACE2) measured alone at 20 nM and mixed at a 1:3 ratio (spike:receptor). (B) Biolayer Interferometry analysis of BwCoV-S binding to beluga ACE2, APN, and DPP4. Responses upon introducing BwCoV-S pre-loaded on NiNTA sensors into recombinant beluga whale proteins ACE2 (a), APN (b), and DPP4 (c) at concentrations of 125 nM (blue), 250 nM (green), or 500 nM (red) recorded on OctetRED384 (Forte Bio). Baseline, Association, and Dissociation phases are separated by dashed line. Association responses were normalised to the average of the 30 s preceding association step.

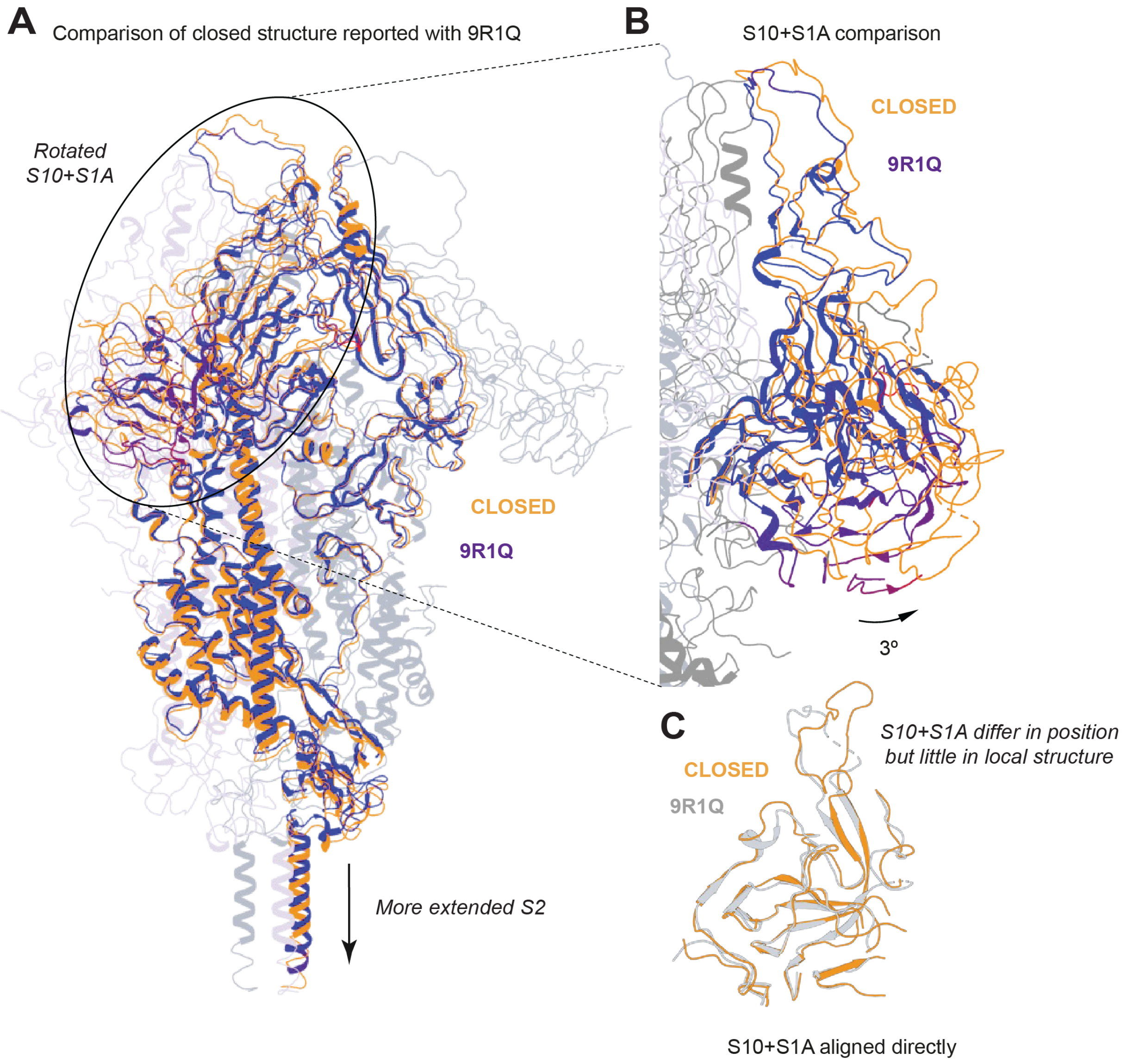


**Figure S4: Comparison of the C3 closed BwCoV-S structure reported here with PDB ID 9R1Q.** Overlays of (A) the whole protomer and (B) the S10-S1A domains of the closed conformation of BwCoV-S reported here and the recently deposited BwCoV-S structure PDB ID: 9R1Q. Our C3-refined structure is used for this comparison because 9R1Q was also reconstructed with C3 symmetry, allowing comparison of the corresponding symmetry-equivalent structures. The reported BwCoV spike in the closed conformation is coloured orange, whilst 9R1Q is coloured by RMSD, where dark blue = 0.07 Å RMSD, dark red = 18.58 Å RMSD, with a mean RMSD of 1.87 Å over the whole protomer aligned on the central helix of S2. (C) Superposition of the S10-S1A domains, with BwCoV-S coloured in orange, and 9R1Q in silver, when aligned on each other, rather than S2.

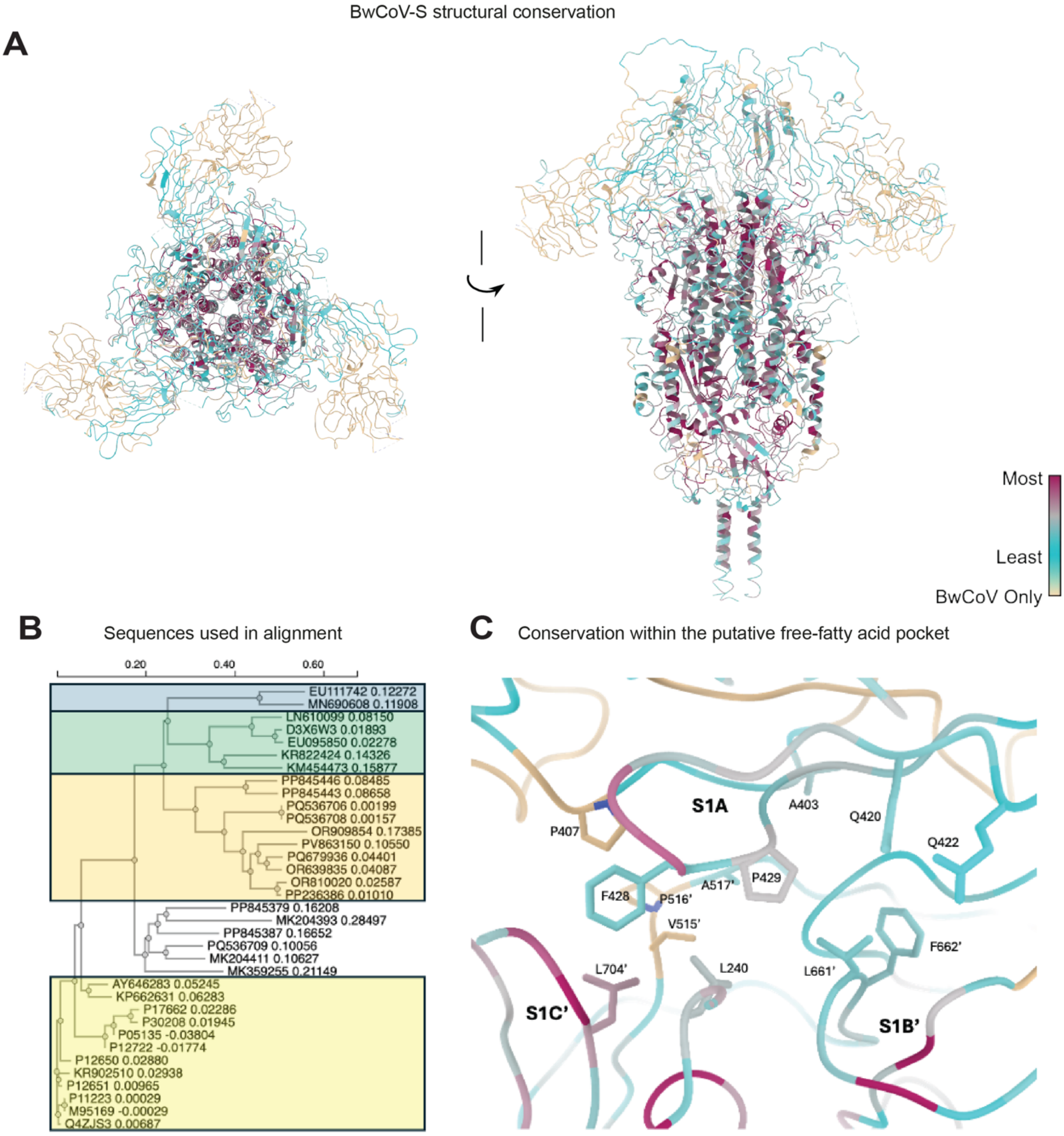


**Figure S5: Conservation of amino acids around the putative free fatty acid pocket.** (A) BwCoV Spike structure in the C3 closed conformation visualising in ChimeraX (with colours representing amino acid sequence conservation as per the inset key) (B) a diverse selection (n=35) of gamma-coronavirus spike protein sequences aligned using the Clustal Omega Multiple sequence alignment tool from EMBL-EBI. (C) Zoom on the putative free fatty acid binding pocket between S1A' and S1B of the adjacent protomer with orientation similar to that in the middle panel of Figure 3.

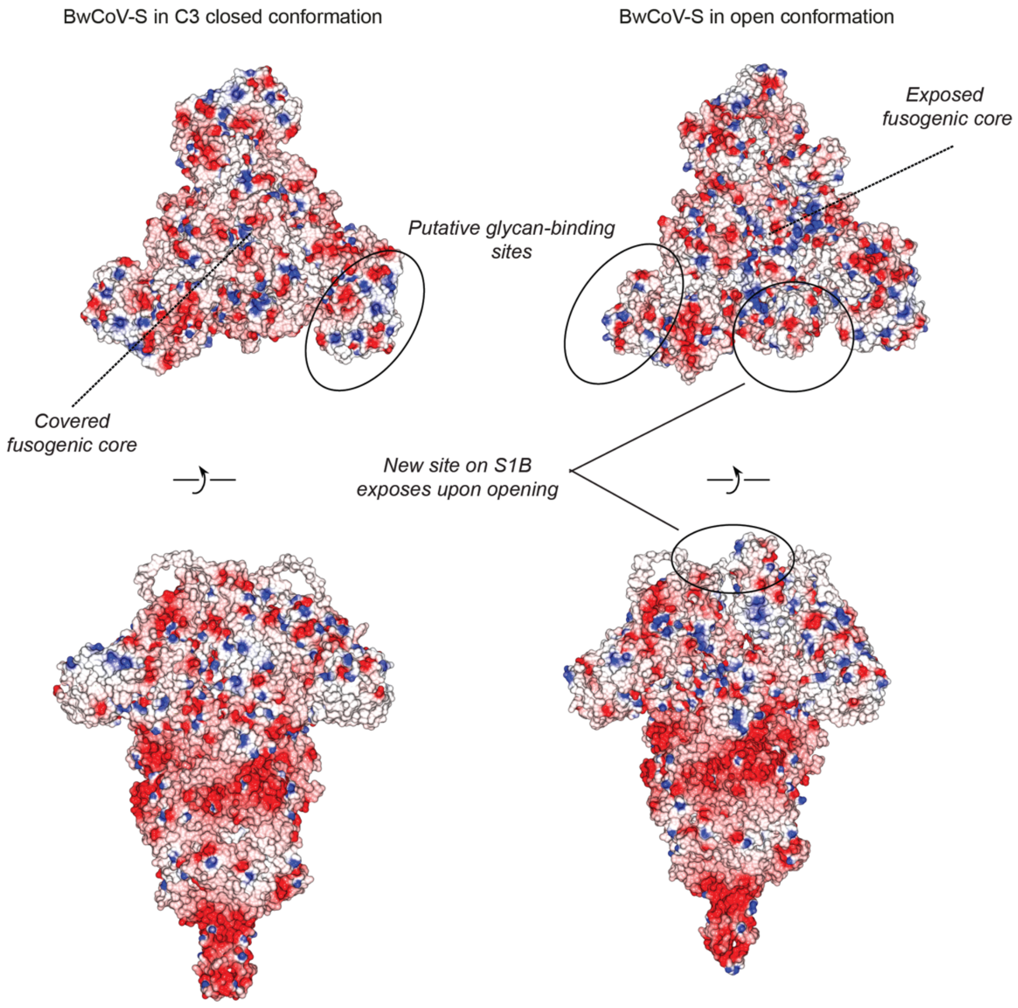


**Figure S6.** Surface electrostatics of BwCoV-S provide clues to potential receptor-binding sites. Surface electrostatic potentials of BwCoV-S in the (left) C3 closed and (right) C1 open conformations, calculated using the built-in Poisson–Boltzmann method in CCP4mg and mapped onto the molecular surface, with negative and positive potentials shown in red and blue, respectively. Structures are viewed from (top panels) the top down, looking towards the virion, and (bottom panels) orthogonal perspectives.

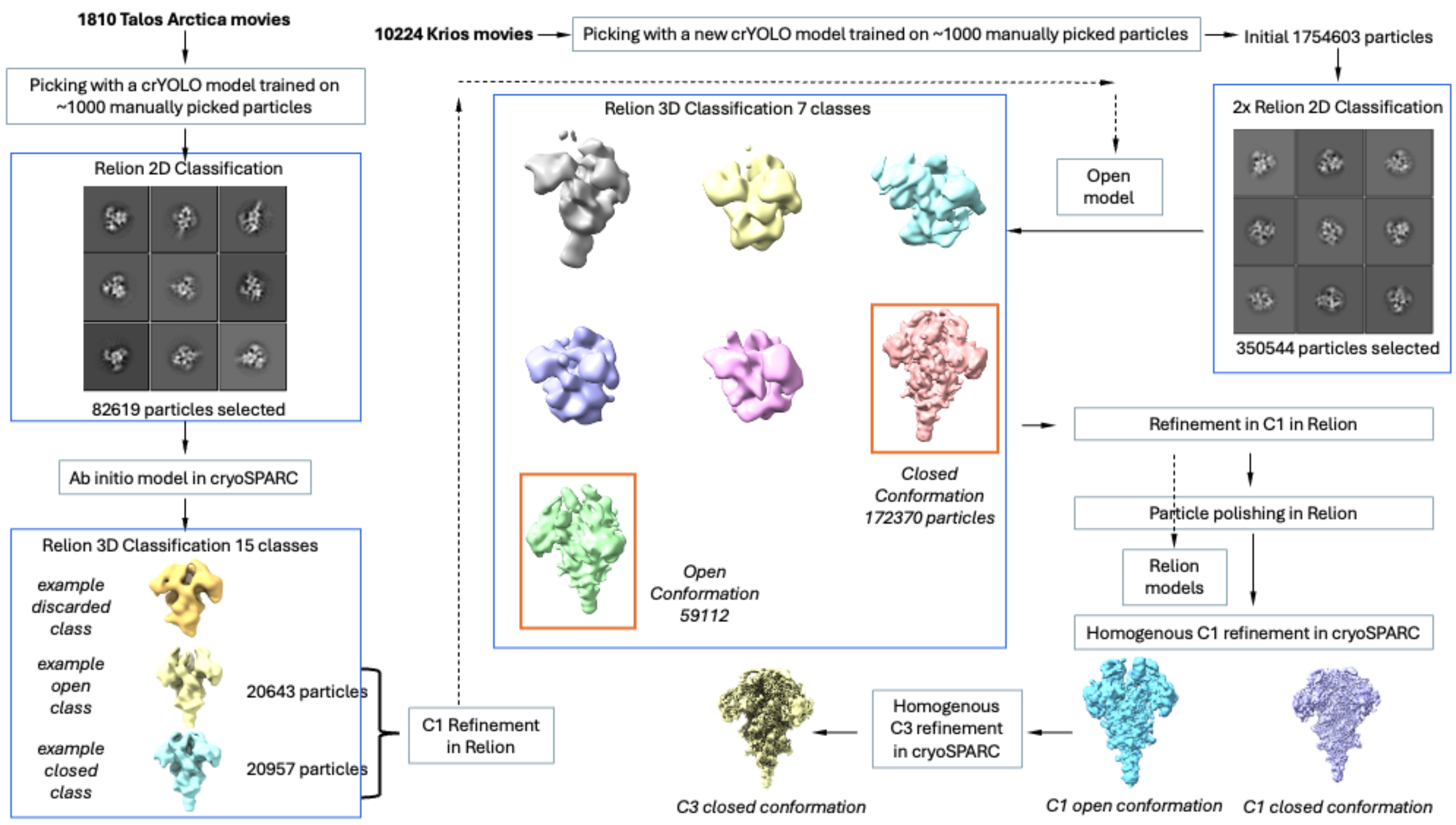


**Figure S7. CryoEM processing schematic.**

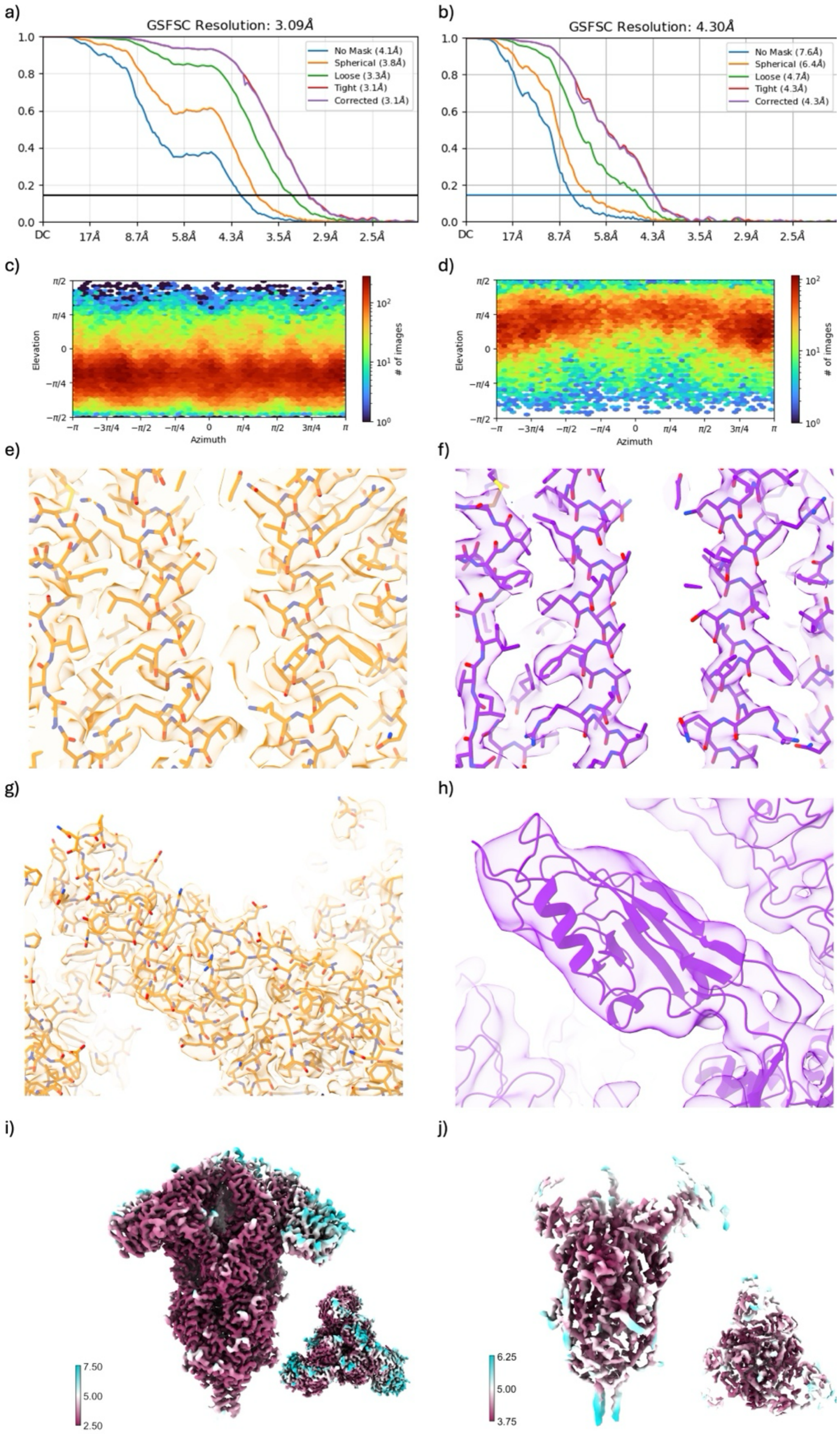


**Figure S8: CryoEM map validation.** Validation of C1 maps corresponding to the closed (panels a, c, e, g, i) and open (b, d, f, h, j). (a,b) Gold Standard Fourier Shell Correlation curves of the final deposited maps and (c,d) particle viewing-direction distribution plots from the final cryoSPARC refinements. Representative map fitting at (e,f) the S2 connecting helix and (g,h) the S1B domain. The maps are displayed at a threshold of 1.000 in UCSF ChimeraX for all panels except (f), wherein 0.350 was used to visualise the positioning of the S1B domain. (g, h) Refined, locally filtered maps coloured by local resolution.

**Table S1: Interactions between domains**

Inter-protomer and intra-protomer analysis of the interfaces of BwCoV-S in C3 closed, C1 closed, and C1 open conformations as identified by PDBePISA. The C3-symmetric closed structure contains equivalent interfaces; the corresponding C3 values are therefore identical. Individual interfaces are reported separately for the asymmetric C1 closed and C1 open structures. Changes upon opening (Δ) were calculated relative to the symmetry-equivalent C3 closed state, as the correspondence between individual protomers of the independently reconstructed asymmetric C1 closed and open states cannot be established. The unprimed, primed, and double-primed protomers correspond to original chains A, B, and C, respectively; original chain A (unprimed) retains the closed S1B orientation in the C1 open structure, while chain B (primed) corresponds to the protomer in which the S10+S1A move away from the symmetry axis in the C1 closed structure.

| Interface | Closed C3 | | Closed C1 | | Open C1 | | Upon opening (open C1 - closed C3) | |
|---|---|---|---|---|---|---|---|---|
| | BSA (Å$^2$) | ΔiG (kcal/mol) | BSA (Å$^2$) | ΔiG (kcal/mol) | BSA (Å$^2$) | ΔiG (kcal/mol) | BSA (Å$^2$) | ΔiG (kcal/mol) |
| **WHOLE PROTOMER INTERFACE** | | | | | | | | |
| Chain:Chain' | 6104.4 | -57.5 | 6104.9 | -58.3 | 4782.9 | -51.8 | -1321.5 | +5.7 |
| Chain':Chain" | 6104.4 | -57.5 | 5705.5 | -53.3 | 4993.4 | -47.8 | -1111.0 | +9.7 |
| Chain":Chain | 6104.4 | -57.5 | 6224.6 | -59.7 | 5126.1 | -63.8 | -978.3 | -6.3 |
| **INTER-PROTOMER** | | | | | | | | |
| S1A:S1B' | 546.6 | -8.6 | 530.5 | -8.2 | 0.0 | 0.0 | -546.6 | +8.6 |
| S1A':S1B" | 546.6 | -8.6 | 91.6 | -0.4 | 2.3 | 0.0 | -544.3 | +8.6 |
| S1A":S1B | 546.6 | -8.6 | 552.0 | -8.9 | 89.2 | -1.5 | -457.4 | +7.1 |
| S1B:S2" | 241.5 | -1.1 | 241.2 | -1.6 | 214.6 | -3.0 | -26.9 | -1.9 |
| S1B':S2 | 241.5 | -1.1 | 230.8 | 0.1 | 0.0 | 0.0 | -241.5 | +1.1 |
| S1B":S2' | 241.5 | -1.1 | 257.4 | -0.9 | 0.0 | 0.0 | -241.5 | +1.1 |
| S1B:S1B' | 127.7 | -1.1 | 129.5 | -0.7 | 0.0 | 0.0 | -127.7 | +1.1 |
| S1B':S1B" | 127.7 | -1.1 | 127.0 | -0.9 | 0.0 | 0.0 | -127.7 | +1.1 |
| S1B":S1B | 127.7 | -1.1 | 131.4 | -0.9 | 0.0 | 0.0 | -127.7 | +1.1 |
| S1A loop:S1B' | 102.7 | 0.1 | 103.4 | -0.7 | 0.0 | 0.0 | -102.7 | -0.1 |
| S1A loop':S1B" | 102.7 | 0.1 | 0.0 | 0.0 | 0.0 | 0.0 | -102.7 | -0.1 |
| S1A loop":S1B | 102.7 | 0.1 | 111.1 | -0.5 | 0.0 | 0.0 | -102.7 | -0.1 |

| Interface | Closed C3 | | Closed C1 | | Open C1 | | Upon opening (open C1 - closed C3) | |
|---|---|---|---|---|---|---|---|---|
| | BSA (Å$^2$) | ΔiG (kcal/mol) | BSA (Å$^2$) | ΔiG (kcal/mol) | BSA (Å$^2$) | ΔiG (kcal/mol) | BSA (Å$^2$) | ΔiG (kcal/mol) |
| S10:S1B' | 30.0 | 0.1 | 20.2 | 0.0 | 0.0 | 0.0 | -30.0 | -0.1 |
| S10':S1B" | 30.0 | 0.1 | 0.0 | 0.0 | 0.0 | 0.0 | -30.0 | -0.1 |
| S10":S1B | 30.0 | 0.1 | 17.3 | -0.1 | 0.0 | 0.0 | -30.0 | -0.1 |
| S1A:S1C' | 564.7 | -6.5 | 573.4 | -7.0 | 567.2 | -7.1 | +2.5 | -0.6 |
| S1A':S1C" | 564.7 | -6.5 | 536.4 | -6.1 | 537.9 | -5.6 | -26.8 | +0.9 |
| S1A":S1C | 564.7 | -6.5 | 587.3 | -6.3 | 531.6 | -6.0 | -33.1 | +0.5 |
| S1D:S2" | 1391.4 | -12.8 | 1353.8 | -15.6 | 1265.6 | -18.8 | -125.8 | -6.0 |
| S1D':S2 | 1391.4 | -12.8 | 1379.1 | -12.4 | 1312.0 | -16.5 | -79.4 | -3.7 |
| S1D":S2' | 1391.4 | -12.8 | 1414.6 | -10.7 | 1317.5 | -13.7 | -73.9 | -0.9 |
| S1C:S2" | 523.9 | -1.6 | 513.3 | -1.3 | 434.9 | -4.8 | -89.0 | -3.2 |
| S1C':S2 | 523.9 | -1.6 | 503.7 | -3.8 | 442.8 | -3.6 | -81.1 | -2.0 |
| S1C":S2' | 523.9 | -1.6 | 509.0 | -3.7 | 471.7 | -3.4 | -52.2 | -1.8 |
| S1A:S2" | 71.1 | -0.1 | 67.3 | -0.6 | 114.2 | -0.3 | +43.1 | -0.2 |
| S1A':S2 | 71.1 | -0.1 | 66.8 | -0.4 | 94.7 | -1.1 | +23.6 | -1.0 |
| S1A":S2' | 71.1 | -0.1 | 73.1 | -0.3 | 93.3 | -1.6 | +22.2 | -1.5 |
| S2:S2' | 2588.4 | -26.6 | 2537.6 | -26.2 | 2302.3 | -21.0 | -286.1 | +5.6 |
| S2':S2" | 2588.4 | -26.6 | 2646.7 | -29.3 | 2501.1 | -22.1 | -87.3 | +4.5 |
| S2":S2 | 2588.4 | -26.6 | 2598.3 | -24.1 | 2438.6 | -29.3 | -149.8 | -2.7 |
| S10:S1C' | 120.4 | -1.6 | 163.8 | -1.6 | 135.9 | -2.2 | +15.5 | -0.6 |
| S10':S1C" | 120.4 | -1.6 | 159.4 | -2.2 | 161.1 | -2.5 | +40.7 | -0.9 |
| S10":S1C | 120.4 | -1.6 | 178.2 | -2.3 | 137.5 | -1.8 | +17.1 | -0.2 |
| **INTRA-PROTOMER** | | | | | | | | |
| S1A loop:S1B | 169.4 | -0.2 | 116.3 | -2.3 | 0.0 | 0.0 | -169.4 | +0.2 |
| S1A loop':S1B' | 169.4 | -0.2 | 0.0 | 0.0 | 0.0 | 0.0 | -169.4 | +0.2 |

| Interface | Closed C3 | | Closed C1 | | Open C1 | | Upon opening (open C1 - closed C3) | |
|---|---|---|---|---|---|---|---|---|
| | BSA (Å$^2$) | ΔiG (kcal/mol) | BSA (Å$^2$) | ΔiG (kcal/mol) | BSA (Å$^2$) | ΔiG (kcal/mol) | BSA (Å$^2$) | ΔiG (kcal/mol) |
| S1A loop":S1B" | 169.4 | -0.2 | 195.5 | -0.3 | 0.0 | 0.0 | -169.4 | +0.2 |
| S1B:S1C | 380.8 | 0.5 | 385.6 | -0.7 | 369.5 | -2.0 | -11.3 | -2.5 |
| S1B':S1C' | 380.8 | 0.5 | 382.0 | -0.6 | 200.9 | -1.0 | -179.9 | -1.5 |
| S1B":S1C" | 380.8 | 0.5 | 384.5 | 0.3 | 212.7 | -1.5 | -168.1 | -2.0 |
| S1A:S1C | 0.0 | 0.0 | 0.0 | 0.0 | 0.0 | 0.0 | 0.0 | 0.0 |
| S1A':S1C' | 0.0 | 0.0 | 0.0 | 0.0 | 0.0 | 0.0 | 0.0 | 0.0 |
| S1A":S1C" | 0.0 | 0.0 | 0.0 | 0.0 | 5.9 | +0.1 | +5.9 | +0.1 |
| S1D:S2 | 340.5 | -5.0 | 372.9 | -5.0 | 292.0 | -3.7 | -48.5 | +1.3 |
| S1D':S2' | 340.5 | -5.0 | 364.8 | -5.2 | 246.5 | -5.0 | -94.0 | 0.0 |
| S1D":S2" | 340.5 | -5.0 | 333.0 | -5.2 | 292.1 | -3.1 | -48.4 | +1.9 |
| S1A:S2 | 68.6 | 0.1 | 65.9 | 0.2 | 96.5 | -0.8 | +27.9 | -0.9 |
| S1A':S2' | 68.6 | 0.1 | 65.2 | 0.1 | 108.9 | -0.2 | +40.3 | -0.3 |
| S1A":S2" | 68.6 | 0.1 | 66.6 | 0.1 | 98.9 | -0.7 | +30.3 | -0.8 |
| S1A:S1D | 433.5 | -4.6 | 416.6 | -6.2 | 419.4 | -4.9 | -14.1 | -0.3 |
| S1A':S1D' | 433.5 | -4.6 | 448.3 | -4.7 | 424.1 | -6.0 | -9.4 | -1.4 |
| S1A":S1D" | 433.5 | -4.6 | 430.3 | -5.1 | 449.9 | -5.5 | +16.4 | -0.9 |
| S10:S1A | 1876.3 | -15.5 | 1899.5 | -16.9 | 1763.3 | -13.4 | -113.0 | +2.1 |
| S10':S1A' | 1876.3 | -15.5 | 1842.4 | -13.0 | 1695.1 | -16.8 | -181.2 | -1.3 |
| S10":S1A" | 1876.3 | -15.5 | 1930.5 | -11.1 | 1568.0 | -14.1 | -308.3 | +1.4 |
| S1C:S1D | 204.3 | -0.5 | 188.6 | 0.0 | 222.3 | -0.4 | +18.0 | +0.1 |
| S1C':S1D' | 204.3 | -0.5 | 203.5 | -0.2 | 208.9 | -0.7 | +4.6 | -0.2 |
| S1C":S1D" | 204.3 | -0.5 | 199.3 | 1.1 | 223.4 | 0.3 | +19.1 | +0.8 |

**Domain definitions:** S10, residues 1–207; S1A, residues 208–354 and 381–476; S1A loop, residues 355–380; S1B, residues 516–668; S1C, residues 502–515 and 669–729; S1D, residues 477–501 and 730–812; S2, residues 813–1331. Contacts between the separately defined S1A loop and the remainder of S1A within the same protomer are not reported because they arise from partitioning the continuous S1A region for PISA analysis.

Abbreviations: BSA, buried surface area; ΔiG, solvation free-energy gain upon interface formation; ΔBSA and ΔΔiG, C1 open minus C3 closed values. Interfaces not identified by PDBePISA in a C1 structure are reported as 0.0 $Å^2$ BSA and 0.0 kcal/mol ΔiG.

**Table S2: Probe list of Glycan Microarray**

| # | Type | Probe* | Structure** |
|---|---|---|---|
| 1 | GAG-related | CS(OS)-DP10-AEAB | 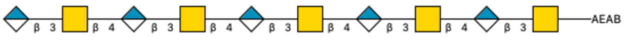 |
| 2 | GAG-related | CS(OS)-DP14-AEAB | 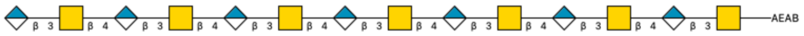 |
| 3 | GAG-related | CSA-DP10(3S)-AEAB | 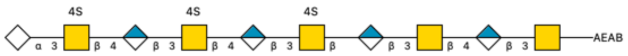 |
| 4 | GAG-related | CSA-DP10(5S)-AEAB | 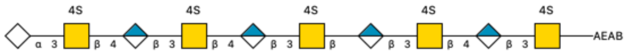 |
| 5 | GAG-related | CSA-DP14-AEAB | 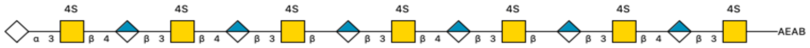 |
| 6 | GAG-related | CSC-DP6-AEAB | 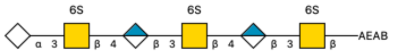 |
| 7 | GAG-related | CSC-DP10-AEAB | 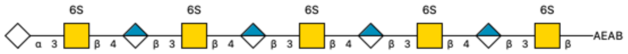 |
| 8 | GAG-related | CSC-DP14-AEAB | 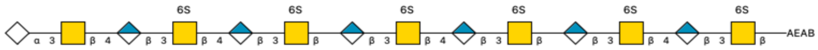 |
| 9 | GAG-related | Hep-DP6-AEAB | 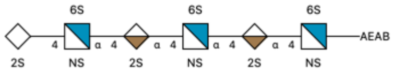 |
| 10 | GAG-related | Hep-DP10-AEAB | 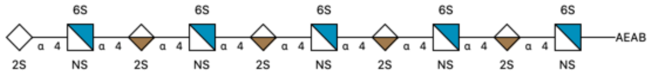 |
| 11 | Blood group + Lewis | H Hexa T1-T2-Lac-PTPA | 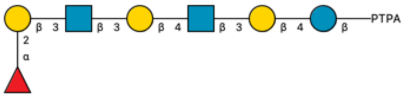 |
| 12 | Blood group + Lewis | H Hexa T2-T1-Lac-PTPA | 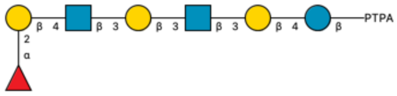 |
| 13 | Blood group + Lewis | A Hexa T1-T2-Lac-PTPA | 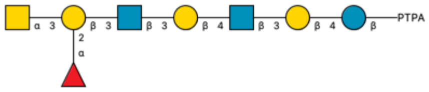 |
| 14 | Blood group + Lewis | A Hexa T2-T1-Lac-PTPA | 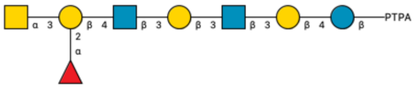 |
| 15 | Blood group + Lewis | B Hexa T1-T2-Lac-PTPA | 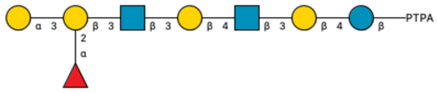 |

| | | | |
|---|---|---|---|
| 16 | Blood group + Lewis | B Hexa T2-T1-Lac-PTPA | 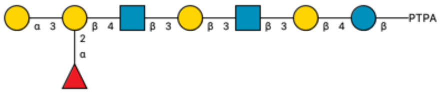 |
| 17 | Blood group + Lewis | Hexa T1(Lea)-T2-Lac-PTPA | 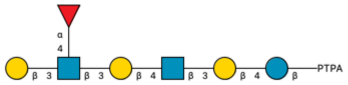 |
| 18 | Blood group + Lewis | Hexa T1(Leb)-T2-Lac-PTPA | 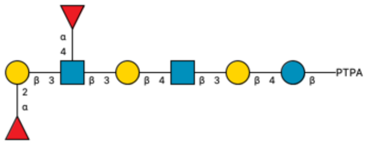 |
| 19 | Blood group + Lewis | Hexa T2(Lex)-T1-Lac-PTPA | 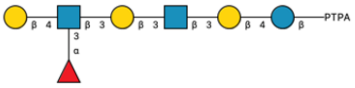 |
| 20 | Blood group + Lewis | Hexa T2(Ley)-T1-Lac-PTPA | 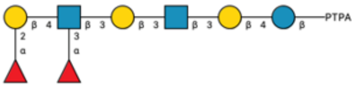 |
| 21 | Gal terminating | LNT-AEAB | 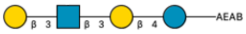 |
| 22 | Gal terminating | LNnT-AEAB | 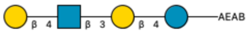 |
| 23 | Gal terminating | Hexa T1-T2-Lac-PTPA | 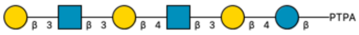 |
| 24 | Gal terminating | Hexa T2-T1-Lac-PTPA | 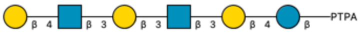 |
| 25 | Gal terminating | Hexa T2-T2-Lac-PTPA | 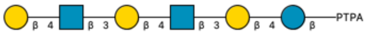 |
| 26 | Gal terminating | KS-DP8(OS)-AEAB | 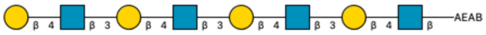 |
| 27 | Gal terminating | Octa T2-T2-T2-Lac-PTPA | 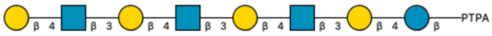 |
| 28 | Mucin cores | Core 1 Thr | 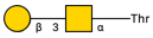 |
| 29 | Mucin cores | Core 2 Thr | 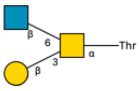 |
| 30 | Mucin cores | Core 3 Ser | 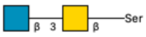 |
| 31 | Mucin cores | Core 4 Thr | 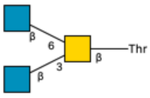 |
| 32 | Sialylated | LSTa-AEAB | 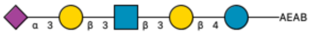 |

| | | | |
|---|---|---|---|
| 33 | Sialylated | LSTd-AEAB | 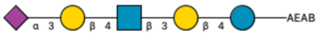 |
| 34 | Sialylated | 3’Sia-Octa-T2-AEAB | 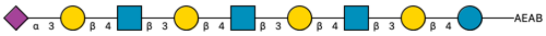 |
| 35 | Sialylated | LSTb-AEAB | 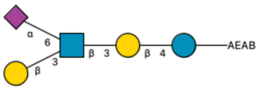 |
| 36 | Sialylated | LSTc-AEAB | 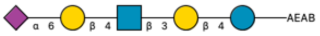 |
| 37 | Sialylated | 6’Sia-Octa-T2-AEAB | 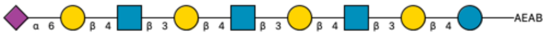 |
| 38 | Sialylated | Fetuin N-Glycan A2 | 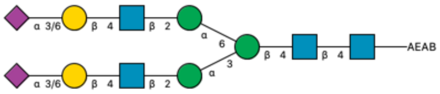 |
| 39 | Sialylated | Fetuin N-Glycan A3-SA3(a) | 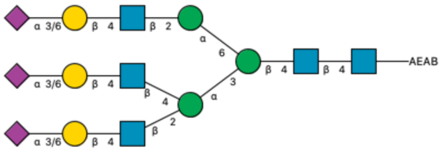 |
| 40 | Sialylated | Fetuin N-Glycan A3-SA3(b) | 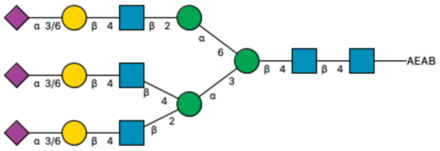 |

** The probe names include an abbreviated description of the glycan moiety followed by the chemical linker abbreviation: AEAB: N-(aminoethyl)-2-aminobenzamide; PTPA: -O-(CH2)3-triazole-(PEG)4-NH2.*

*** The glycan probe structures are depicted using Symbol Nomenclature for Glycans (SNFG; https://www.ncbi.nlm.nih.gov/glycans/snfg.html).*

**Table S3: Model refinement statistics**

| Parameter | Closed C3 | Closed C1 | Open C1 |
|---|---|---|---|
| Map resolution (GSFSC = 0.143; Å) | 2.82 | 3.09 | 4.30 |
| D FSC Model (0.143) | 2.7 | 2.9 | 4.0 |
| $CC_{1/2}$ Mask | 0.83 | 0.84 | 0.80 |
| No. Atoms | 30198 | 30198 | 30198 |
| B-factors (Å$^2$) min/max/mean | 14.01 / 147.08 / 49.32 | 23.49 / 203.97 / 65.49 | 28.04 / 408.51 / 170.94 |
| Bond Length RMSD Deviations (Å) | 0.003 | 0.007 | 0.005 |
| Bond Angle RMSD Deviations (°) | 0.524 | 0.823 | 1.299 |
| Clash Score | 3.59 | 6.91 | 17.50 |
| Ramachandran (%) Favoured/Allowed/Outliers | 96.44 / 3.41 / 0.16 | 94.05 / 5.82 / 0.13 | 91.80 / 7.81 / 0.39 |
| Rotamer Outliers (%) | 0.06 | 0.15 | 0.06 |
